# A polinton-like virus of the algae *Chrysochromulina parva* inhibits the growth of a newly isolated relative of *Tethysvirus ontarioense*

**DOI:** 10.64898/2026.01.07.698126

**Authors:** G. Thomas, I. Inamoto, C. Palermo, G. Bajaj, S. M. Short

**Affiliations:** University of Toronto Mississauga 3359 Mississauga Road, Mississauga, Ontario, Canada

**Keywords:** Algal virus, virophage, haptophyte, Nucleocytoviricota, Preplasmiviricota, Mesomimiviridae, Phycodnaviridae, CpV, CpV-PLV Moe

## Abstract

Previous discovery of genomes of polinton-like viruses (PLVs) associated with viruses of *Chrysochromulina parva* stimulated this research to determine the biological nature of these putative viral hyperparasites. Purification of *C. parva* viruses to enable co-infection experiments led to the discovery of a previously unknown virus, CpV-BQ3, which, based on sequence information and electron microscopy, is a species of *Tethysvirus*, a genus within the *Megaviricetes*. Purification and TEM imaging of CpV-PLV Moe revealed naked icosahedral particles morphologically similar other cultivated virophages and PLVs. Mixed-infection experiments with the putative polinton-like virus CpV-PLV Moe demonstrated that CpV-BQ3 supports its replication whereas the putative Phycodnavirus CpV-BQ1 does not. Further, experimental infections with differing proportions of the Moe and its helper virus CpV-BQ3 revealed a dose-effect whereby high levels of Moe had a greater negative impact on BQ3 replication compared to lower levels. Conversely, high levels of Moe relative to BQ3 provided greater protection for *C. parva* allowing enhanced cell survival whereas low doses of Moe did not prevent cell lysis. Overall, the results of this study demonstrated the intimate relationship of CpV-PLV Moe with the newly discovered virus, CpV-BQ3, and *C. parva*, and illustrate the complex ecology of algal viruses.

## 1. Introduction

The discovery of Sputnik [1], a satellite virus that coinfects the amoeba *Acanthamoeba castellanii* with a giant dsDNA virus, Acanthamoeba castellanii mamavirus Hal-V (*Mimivirus bradfordmassiliense*), marked the beginning of research on virophage biology and their relationship to other viruses and mobile genetic elements (MGEs). The observation that Sputnik depended on mamavirus as a helper virus while also compromising mamavirus’ replication led the researchers to introduce the term *virophage* to highlight its hyperparasitic lifestyle and allegorically relate it to bacteriophage [1]. Soon after, Mavirus, a relative of Sputnik that coinfects the marine phagotrophic flagellate *Cafeteria sp.* with its helper virus, CroV, was described as a virophage with evolutionary connections MGEs in the Polinton, or Maverick, family of large DNA transposons of eukaryotes [2]. Around the same time, virophage genomes were reported from Organic Lake Antarctica metagenomes [3], and numerous examples of complete or near-complete virophage and Polinton-like virus (PLV) genomes have now been observed in metagenomes collected from around the world [4–8]. Isolation of Zamilon (*Sputnikvirus zamilonense*) provided evidence for the variable replication strategies of virophages because it was specific to ‘group C’ *Mimiviridae* (i.e., *Megavirus chilense*) and did not appear to have a significant impact on the replication of its helper virus [9]. Together, these and related literature triggered debate about the nature and classification of virophages [e.g., 10, 11-13], their shared evolutionary history [e.g., 14, 15, 16], and their relationship to vertebrate adenoviruses and prokaryote tectiviruses like the bacteriophage PRDI [17].

Virophages, PLVs, adenoviruses, and tectiviruses are currently classified in the phylum *Preplasmiviricota* and are placed in the kingdom *Bamfordvirae* along with dsDNA viruses of eukaryotes in the phylum *Nucleocytoviricota*, as defined by the International Committee on Taxonomy of Viruses (ICTV); *ICTV Virus Taxonomy* https://ictv.global/taxonomy [18]. Given the rapid pace of discovery in this area of virology, it is appropriate that virophage taxonomy has evolved since their discovery. For example, the genera *Mavirus* and *Sputnikvirus* were previously classified in the family *Lavidaviridae* [19], but they are now separated into different orders with *Sputnikvirus* as a member of the *Mividavirales* and *Mavirus* in the order *Lavidavirales* [13]. Even the use of the term ‘virophage’ has varied in the literature. It has recently been argued that the term virophage can be used in an informal sense to describe dsDNA viruses that engages in a hyperparasitic replication strategy by parasitizing dsDNA viruses of the phylum *Nucleocytoviricota*, yet the term can also be used in a formal sense to refer to viruses in the class *Virophaviricetes* in the phylum *Preplasmivircota* [17]. Since the time of their discovery, it has become ever more evident that the biology and natural history of preplasmiviricotes is both fascinating and complex, bridging the evolution of eukaryote and bacterial viruses and providing a direct link between mobile genetic elements and viruses.

Related to the virophages (*sensu stricto*) are the Polinton-like viruses (PLVs) which resemble polintons with respect to their size and genetic composition but are distinct because they lack genes such as the retrovirus-like integrase, DNA polymerase B, or maturation protease typically found in polintons. However, like polintons, PLVs encode the hallmark double jelly-roll major capsid proteins (MCP) of diverse dsDNA viruses in the *Nucleocytoviricota* [14, 20]. Genomic analysis of PgV-16T, a virus of the marine haptophyte (Prymnesiophyceae) *Phaeocystis globosa*, led to the assembly of its ∼460 kbp genome as well as a smaller ∼20 kbp linear genome labelled PgVV because it was, at the time, considered virophage-like [21]. Subsequent research demonstrated that PgVV was distinct from characterized Polintons and virophages, and searches for PgVV-like MCP genes in genomic and metagenomics sequences led to the identification of a novel group of PLVs [5]. Like the discovery of virophages in metagenomes from around the world, subsequent studies of PLV diversity in environmental samples and metagenomes led to the identification of more than 80 PLVs from a single alpine lake in Austria and over 550 PLVs in metagenomes and eukaryotic genomes [22]. Additionally, endogenous viral elements, or EVEs, ranging from 14 to 40 kbp and related to virophages and PLVs have been identified in diverse protist genomes spanning the breadth of eukaryote clades [23], and PLVs have even been observed recently as MGEs integrated into a broad range of stony coral (order Scleractinia) genomes [24]. Further, exploration of virus genes in genome assemblies of *Tetraselmis* spp., a genus of green algae, demonstrated that close relatives of Tsv-N1, an unclassified virus of *T. striata* that does not require a helper virus [25], were integrated into the genomes of many green algae highlighting the fact that these viruses exist as both extracellular virions and integrated proviral elements [26]. Recognizing the distinct identity of PLVs, the phylum *Preplasmiviricota* has been reorganized to include the established classes *Polintoviricetes* and *Maveriviricetes* (now named *Virophaviricetes*), as well two new classes *Pharingeaviricetes* and *Aquintoviricetes. Pharingeaviricetes* is the home for adenoviruses whereas *Aquintoviricetes* is the class within which the algal-infecting PLVs TvV-S1 [a virus presumably related to Tsv-N1; 5, 27] and PgVV are placed [17].

The PLV PgVV is particularly important to the study described here. Viruses of the haptophyte alga *Phaeocystis globosa* (PgVs) were isolated before mimiviruses, and at that time known algal viruses were all classified as members of the class *Phycodnaviridae*. Several, but not all, of the PgV isolates were most closely related to CbV-PW1, viruses isolated from the marine prymnesiophyte *Chrysochromulina brevifilum* [28] and thus were tentatively considered prymnesioviruses [29]. Further characterization of the viruses infecting *P. globosa* revealed that there were in fact 2 distinct groups of viruses that could be discriminated based on the sizes of their virions and genomes [30]. The smaller viruses (Group II PgVs) encode DNA polymerase genes most closely related to the phycodnavirus CbV-PW1 whereas the larger viruses (Group 1 PgVs) are now known to be related to mimiviruses. Following the discovery and rapid expansion of viruses related to *Mimivirus*, it became clear that many giant viruses of algae were more closely related to mimiviruses than phycodnaviruses, and for a time these viruses were loosely classified as ‘extended mimiviridae’ [31]. These giant viruses of algae are now classified in the order *Imitervirales* family *Mesomimividae*, and the *P. globosa* virus PgV-16T has been named *Tethysvirus hollandense* [32, 33]. PgVV, The PLV identified in through genomic analysis of PgV-16T, has not yet been isolated, but a recent report described the isolation and life history of another very closely related PLV that coinfects *P. globosa* along with PgV-14T, a strain of *T. hollandense*. This PLV, called Gezel-14T, has been confirmed as bona fide virus with a virophage-like lifestyle; although it has a fitness impact on its helper virus, it doesn’t appear to protect the host cell against infection [34]. Gezel-14T is considered a member of *Aquintoviricetes*.

A strikingly similar system of viruses and PLVs have been observed in association with the freshwater haptophyte *Chrysochromulina parva* [35–37]. *C. parva* itself has a challening taxonomic history with recent evidence that the isolate *C. parva* CCMP291 may be mislabled and should be classified as *C. Tobinii* [38, 39]. Nonetheless, this species of haptophyte, like *P. globosa*, is infected by viruses in the orders *Algavirales* and *Imitervirales*, and is also associated with PLVs. When originally characterized, viruses infecting *C. parva* CCMP291 were named *Chrysochromulina parva* virus BQ1 (CpV-BQ1) and were putatively classified as a phycodnaviruses based on phylogenetic analysis of cloned PolB gene fragments [37]. Shortly thereafter, high-throughput sequencing resulted in assembly of a ∼430 kbp virus genome encoding 503 ORFs with a PolB gene distinct from BQ1, and this virus was labelled CpV-BQ2 [36]. CpV-BQ2 is now formally classified as *Tethysvirus ontarioense*, a mesomimivirus whose closest relatives are group I PgVs [32]. Like *P. globosa*, *C. parva* CCMP291 may also be subject to tripartite or hyperparasitic infections because ∼23 kbp genome sequences of three PLVs, CpV-PLV Larry, Curly, and Moe, were also identified during assembly of the BQ2 genome. The striking similarities of the infections of *C. parva* (*C. tobinii*) and *P. globosa* allow the speculation that this tripartite infection system with a haptophyte host is derived from a common ancestor of these algae and predates their transition to freshwater environments. Whatever their evolutionary history, the life history and impacts on host survival and viral replication of the PLVs of *C. parva* CCMP291 have not been studied. In this study, we show that CpV-PLV Moe is a genuine virus with a virophage-like replication strategy that depends on co-infections with a newly discovered strain *of Tethysvirus ontarioense* which we have called CpV-BQ3.

## 2. Materials and Methods

### 2.1 Cell and virus cultivation

Non-axenic cultures of *Chrysochromulina parva* CCMP291 were originally obtained from the National Center for Marine Algae and Microbiota (NCMA; https://ncma.bigelow.org/) and have been maintained in the laboratory for over a decade following NCMA’s culturing recommendations. Briefly, naïve cells that haven’t been exposed to any lytic agents were maintained in 100 to 150 mL batch cultures grown in DY-V medium [40] at a temperature of 15 °C with a 12: 12 h light: dark cycle at a photon flux density of 23 μmol m⁻² s⁻¹. When cells reached a maximum density of approximately 3.0 x 10^6^ cells mL^-1^, after roughly three weeks of growth, they were diluted approximately 100-fold by aseptically transferring a small aliquot of the culture into fresh medium.

Viral lysates have also been maintained in the lab for over a decade since they were first reported [37]. Following addition of 0.5 to 1 mL of undefined, filtered lysates into a mid-log phase culture (i.e., *C. parva* abundance ranging from 0.3 to 1.0 cells mL^-1^), viruses were filter-purified after cell lysis as characterized by a loss of pigmentation in the culture (i.e., cultures clear), or when cell concentrations decreased below 3 × 10⁵ cells mL^-1^. Filter purification involved sequentially vacuum filtering lysates through 47 mm diameter, 0.50 μm nominal pore-size borosilicate glass microfiber filters (GC50, Advantec) and 47 mm diameter 0.45 μm pore-size PVDF Durapore® membranes (MilliporeSigma). Additionally, to further purify some virus preparations (see below), lysate solutions were also filtered through a 0.1 μm pore-size Minisart® (Sartorius), or 0.20 μm pore-size Filtropur S (Sarstedt) sterile syringe filters. Following filtration, lysate stocks were stored in the dark at 4 °C for up to 7 months.

### 2.2 Virus isolation and cultivation

To isolate individual virus types from mixed lysates, a combination of serial propagation, end-point dilution and differential filtration was used, and the abundances of known viruses (i.e., CpV-BQ1 and - BQ2, and CpV-PLV Larry, Curly, and Moe) were monitored in all lysate samples using qPCR as described below. Every lysis experiment conducted throughout this study was initiated from a single culture stock which was split to include independent mock-infected cultures (i.e., as a cell growth control) and cultures infected with putative lytic agents. Starting with a mixed viral stock that included all known viruses, viruses were serially propagated in two parallel series by adding 0.5 mL of 0.45 or 0.20 μm pore-size syringe filtered lysate into a mid-log phase *C. parva* culture and then filtering and transferring the resulting lysate into fresh mid-log phase cultures. After five generations, a lysate from each filtration series was subjected to end-point dilution in 96-well microtiter plates by inoculating 8 wells containing 250 μL of mid-log cells with 50 μL of 10-fold serially diluted lysates. In each plate, 16 wells were used as growth controls and the dilution factors ranged from 10^0^ to 10^9^. The entire 300 μL sample from a well which cleared (i.e., cells lysed) at the highest dilution level, or lowest concentration of lytic agents, were inoculated into 100 mL cultures to scale up the lytic agents, and these were serially propagated again for several generations of replication. Finally, a portion of these scaled-up lysates were filtered through 0.1 μm pore-size syringe filters to determine if this filtration step eliminated any observable lytic activity.

### 2.3 Sequence Analysis

An infectious *C. parva* lysate containing no detectable levels of CpV-BQ1 or -BQ2, or CpV-PLV Larry, Curly, or Moe was concentrated by centrifugation at 31,000 rpm for 1 hour at 20°C using an SW32Ti rotor (Beckman-Coulter). The supernatant was then discarded by inverting the centrifuge tube and residual liquid was used to resuspend any pelleted material. DNA was extracted from 300 µL of this resuspended material using a Promega Maxwell RSC Viral Total Nucleic Acid extraction kit. Final DNA concentrations were measured using a Qubit dsDNA HS assay kit (Thermo Fisher Scientific). Sample library preparation was performed using an Illumina DNA Prep kit and UDI indices and libraries were sequenced from paired ends (2 x 151 bp) on a NextSeq 2000 (Illumina) by SeqCenter (Pittsburgh, PA). Demultiplexing, quality control, and adapter removal were performed by SeqCenter using BCL-convert version 3.9.3 (Illumina).

Additional quality control was performed using Sickle version 1.33 [41] with a quality threshold value of 30 and read length cut-off value of 50. Reads were assembled using default parameters in metaSPAdes version 3.15.5 (https://www.ncbi.nlm.nih.gov/pmc/articles/PMC5411777/). BLASTx searches against the October 2022 NCBI-nr database (https://ftp.ncbi.nlm.nih.gov/blast/db/FASTA/nr.gz) were performed using DIAMOND version 2.0.15 [42] with frameshift alignment and more sensitive modes activated. MEGAN6-LR version 6.24.1 [43] was used to annotate contigs using the February 2022 protein accession mapping file (https://software-ab.informatik.uni-tuebingen.de/download/megan6/welcome.html) with long-read mode activated and bit score and E value cutoffs of 100 and 10^−6^, respectively. The inferred amino acid sequences of annotated genes from Illumina sequencing were compared to GenBank NR using BLASTp. Homologous sequences of common *Nucleocytoviricota* marker genes encoding PolB (DNA polymerase family B), A32 (packaging ATPase), and VLTF3 (Poxvirus Late Transcription Factor) proteins [e.g., 33] were analyzed phylogenetically using MEGA version 12 [44]. For all phylogenetic analyses, the percentage of replicate trees in which the associated taxa clustered together (500 replicates) is shown next to the branches [45]. The initial trees for the heuristic searches were selected by choosing the trees with the superior log-likelihood between Neighbor-Joining [46] and Maximum Parsimony trees. The NJ trees was generated using a matrix of pairwise distances computed using the Jones-Taylor-Thornton model [47], whereas the MP trees had the shortest lengths among 10 MP tree searches, each performed with a randomly generated starting tree.

### 2.4 Transmission Electron Microscopy

Putatively purified virus stocks were analyzed using transmission electron microscopy. Prior to analysis, samples containing no detectable viruses yet retaining lytic activity, or samples containing high concentrations of the polinton like virus (PLV), CpV-PLV Moe, as inferred from qPCR, were concentrated by centrifugation at 33,000 RPM for 1 h for the unknown lytic agent, or 11,000 RPM for 1 h for the CpV-PLV Moe sample using the SW32Ti Rotor in an Optima L-80XP Ultracentrifuge (Beckman-Coulter). Pellets were resuspended in sterile 10 mM Tris-Cl, pH 8.5. Carbon-coated copper grids (FCF400-Cu-UB, Electron Microscopy Sciences) were glow discharged immediately before use, and 10 µL of concentrated samples were applied to the grid and were allowed to sit for 10 minutes. The grid was then washed with double distilled H_2_O three times followed by staining using 0.5 to 2% uranyl acetate for 30 seconds. Stained grids were visualized using a Talos L120C transmission electron microscope (Thermo Fisher Scientific) at the Microscopy Imaging Laboratory, Temerty Faculty of Medicine, University of Toronto.

### 2.5 Cell counts, MPN, and qPCR

Cell counts were performed on culture samples by adding 20 μL of Lugol’s iodine to 200 μL samples immediately after collection. Samples were stored in the dark at room temperature and were counted within 2 weeks following collection using a Bright-Line hemocytometer (Hausser Scientific) and a ZEISS Primostar 3 microscope equipped with an Axiocam 105 digital camera (ZEISS). Images of five of the nine grid blocks of the hemocytometer were captured, and the images were processed in Fiji [48], where a pseudo–flat field correction (blurring parameter = 50) was applied to even out background illumination, followed by thresholding to highlight cells for counting. Raw counts were converted to estimates of cell abundance following standard hemocytometer instructions.

The infectivity of stocks of the viruses CpV-BQ3 or CpV-BQ1 were estimated using the Most Probable Number (MPN) assay. Eleven 10-fold serial dilutions of CpV-BQ3 or CpV-BQ1 lysates were prepared in sterile DY-V medium, and 50 μL of each virus dilution were inoculated into 16 wells of a 96-well microtiter plate containing 250 μL of a *C. parva* culture grown to mid-log phase (see above). In each plate, two columns of wells (i.e., 16 wells) were mock-infected with 50 μL of sterile DY-V media (16 wells) and served as controls for *C. parva* growth and as comparators to assess lysis based on color changes. All MPN plates were incubated in the same growth conditions used to maintain *C. parva* cultures (as detailed above). Plates were checked and scored for lysis (i.e., well clearing) regularly up to 21 days to ensure that lysis patterns had stabilized with no other clearing. The number of wells lysed at each dilution level was used to estimate the infectivity of virus stocks of CpV-BQ3 or CpV-BQ1 with the Excel-based MPN calculator MPN_ver6 [49]. Because infection with CpV-PLV Moe alone did not lyse *C. parva* cultures, infectivity estimation via MPN was not possible. Therefore, the abundance of CpV-PLV Moe was only estimated using qPCR.

Gene copy abundances were used to infer virion abundance using the 5′ nuclease assay (i.e., TaqMan® based quantitative PCR) targeting PolB genes for CpV-BQ1, -BQ2, and -BQ3; and the major capsid protein (MCP) genes for CpV-PLV Larry, Curly, and Moe (Table 1). Quantitative PCR (qPCR) was conducted as previously described for CpV-BQ1 and other freshwater viruses [37, 50]. Briefly, 25 μL reactions contained 1 X PCR buffer [20 mM Tris-HCl (pH 8.4) and 50 mM KCl], 5.0 mM MgCl_2_, 200 μM each dNTP, 400 nM each forward and reverse primer, 200 nM of TaqMan® probe, 0.625 U Invitrogen Platinum® Taq DNA polymerase (Thermo Fisher Scientific), 1 X ROX Reference Dye (Life Technologies), and 2 μL of virus sample. Virus samples were used directly in the qPCR reaction after storage at -20 °C for at least 15 h. Reactions were cycled in a Stratagene MX3000P thermal cycler (Agilent Technologies) with the following parameters: 95°C for 5 min, and 40 cycles of 95°C for 15 s followed by 60°C for 1 min. Primer and probe sets not previously reported (Table 1) were designed and validated as previously described [50, 51]. To determine if gene-specific primers and probes performed with acceptable specificity, each primer and probe set used in this study was tested in amplifications of PCR-derived gene fragments cloned into pGEM®-T vector plasmids, which also served as the standards for all qPCR assays as described previously [50].

**Table 1.**
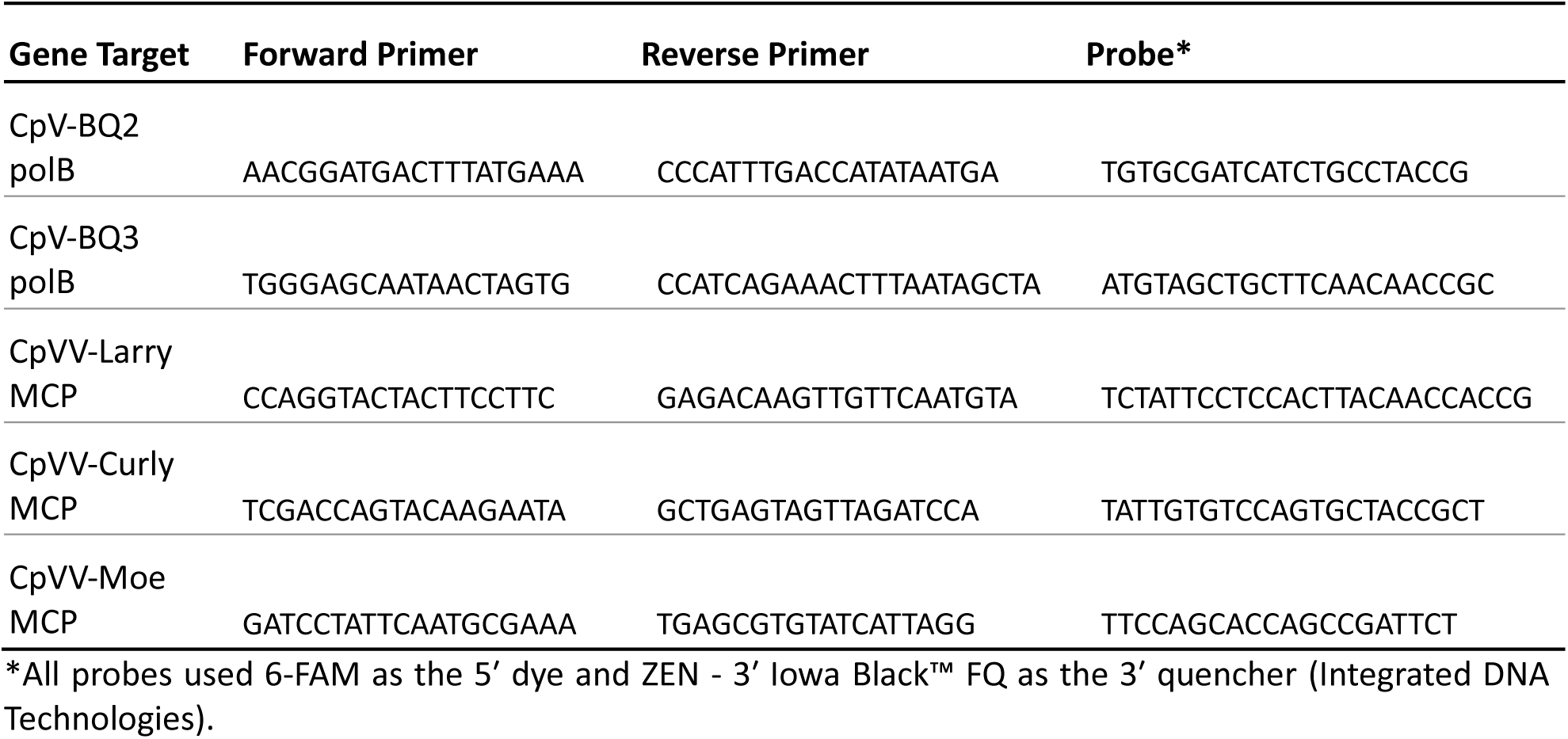
Oligonucleotides used for qPCR (shown in 5′ to 3′ orientation)

### 2.6 Infection experiments

The infectivity of CpV-BQ3 stocks were quantified using MPN, whereas virion concentrations were estimated for CpV-PLV Moe using qPCR. Following cell and virus quantification, each experiment trial was set up with four treatments established in triplicate using disposable 75 cm², 250 mL capacity, vented, Polystyrene cell culture flasks (Sarstedt) for suspension cultures. For each experiment, the starting abundance of *C. parva* cells were approximately 3.5 x 10⁵ cells mL^-1^. Each treatment involved inoculating stocks of CpVs and/or CpV-PLV Moe into 100 mL of *C. parva* cultures. Inoculations never exceeded 1 mL in total and were assumed to have no impact on the composition of the growth medium and only a minimal impact on cell concentrations. Each experimental trial included a no-infection control with *C. parva* growing on its own, as well as a lytic infection control with *C. parva* inoculated with CpV-BQ3 at MOI of 0.01 infectious units per cell, as determined via MPN assays; the same MOI was used for all experimental infections involving CpV-BQ3. Treatment groups included co-inoculation of *C. parva* with CpV-BQ3 and CpV-PLV Moe at proportions as determined via qPCR. Specifically, the experimental treatments included mixed infections at gene copy ratios of BQ3: Moe at 1:10, 1:200, 1:500, 1:900, and 1:1000. Additional treatments consisted of CpVV-Moe-only infections and co-infections of CpV-BQ1 with CpV-PLV Moe at 1:500. The concentration of the CpV-PLV Moe inoculum used in the ‘Moe alone’ treatment was comparable to the gene copy concentration of the CpV-PLV Moe inoculum in the 1:1000 co-infection treatment. The CpV-BQ1 inoculum was matched to the gene copy number of the CpV-BQ3 inoculum used in its respective infection control.

Treatments were distributed across four separate experimental trials for logistical reasons related to the size of the growth chamber and minimize the time required for sample collection. To permit comparisons across independent experimental trials, and as stated above, each trial included a no-infection control, an infection control, and two experimental treatments except for the last trial which included only one treatment; all treatments and controls were conducted in triplicate in independent culture flasks. The experimental treatments included in each trial were: 1) BQ3+Moe 1:1000 and Moe alone; 2) BQ3+Moe 1:900 and BQ3+Moe 1:10; 3) BQ3+Moe 1:500 and BQ1+Moe 1:500; and 4) BQ3+Moe 1:200. Thus, trials 1-3 consisted of twelve culture flasks in total whereas trial 4 included only nine.

All flasks which were incubated in a growth chamber at 15 °C under a 12:12 h light–dark cycle for a total duration of 20 days. Flasks were arranged within the growth chamber to maximize variability of irradiance for each replicate flasks across the averaged irradiance of approximately 23 μmol photons m⁻² s⁻¹. Sampling was performed consistently at the same time each day. Prior to sampling, flasks were gently inverted five times to homogenize the cultures. During the first seven days, two 200 μL samples were aseptically collected every 24 h from each flask for cell counts and qPCR. Subsequently, sampling frequency was reduced to every 48 h on Days 9, 11, 13, and 15. After Day 15 sampling was limited to one final timepoint on Day 20. As noted above, samples collected for cell counts were enumerated with 2 weeks following collection, whereas samples collected for qPCR were stored at -20 °C immediately upon collection and until further analysis; qPCR analysis was conducted within 3 weeks following sample collection.

### 2.7 Statistical Analyses

For each experimental run, *C. parva* cell abundances on day 9 were compared among treatments using Welch’s ANOVA, followed by Games–Howell post hoc testing. Data were power-transformed prior to analysis to improve normality, and the transformed residuals were assumed to follow an approximately normal distribution based on Q–Q plots. All post hoc tests were performed using the rstatix package in R.

To evaluate whether the CpV-PLV Moe affected CpV-BQ3 replication, log₁₀-transformed BQ3 abundances from day 9 samples were compared between the BQ3-only infection controls and corresponding BQ3+Moe co-infections using a Welch’s two-sample t-test. A normal distribution of the log-transformed data was assumed. To conservatively account for multiple comparisons, a Bonferroni-adjusted significance threshold of α = 0.025 was applied to all tests.

Reductions of CpV-BQ3’s replication (i.e., BQ3 abundance in co-infections expressed as a fraction of the mean BQ3-only abundance within each experimental run) were compared across treatments on Day 9 using Welch’s ANOVA with Games–Howell post hoc testing. Reduction values were power-transformed prior to analysis to improve normality, and the transformed residuals were assumed to follow an approximately normal distribution based on Q–Q plots. Post hoc tests were performed using rstatix in R.

## 3. Results

For the sake of brevity, the viruses CpV-BQ1, -BQ2, and -BQ3 will henceforth be referred to as simply BQ1, BQ2, and BQ3. Likewise, the PLVs CpV-PLV Larry, CpV-PLV Curly, and CpV-PLV Moe will be referred to as Larry, Curly, and Moe.

### 3.1 Quantitative PCR assay validation

Each primer and probe set used in this study was tested in amplification reactions with cloned gene fragments of each virus and PLV when added to the reaction at a concentration of 5.0 x 10^9^ gene copies mL^-1^ (Table 2). Considering the Cq value as an indication of amplifiability of different template molecules, the primer and probe set targeting Curly was the most prone to off-target amplification producing a Cq value of 20.3 when amplifying Moe’s MCP gene compared to a Cq of 15.0 for its intended target template. On the other hand, the primer and probe set targeting Moe was the most specific producing no amplification for any target except Curly; in this case, the on-target Cq was 17.2 whereas the off-target Cq was 36.0. Similarly, the primer and probe set targeting the newly discovered BQ3 virus (see below) produced no amplification for any templates except BQ2 and Moe. For this primer and probe set, the on-target Cq was 17.2 whereas the off-target Cq values were 27.0 and 36.8 for the templates of BQ2 and Moe, respectively.

**Table 2.**
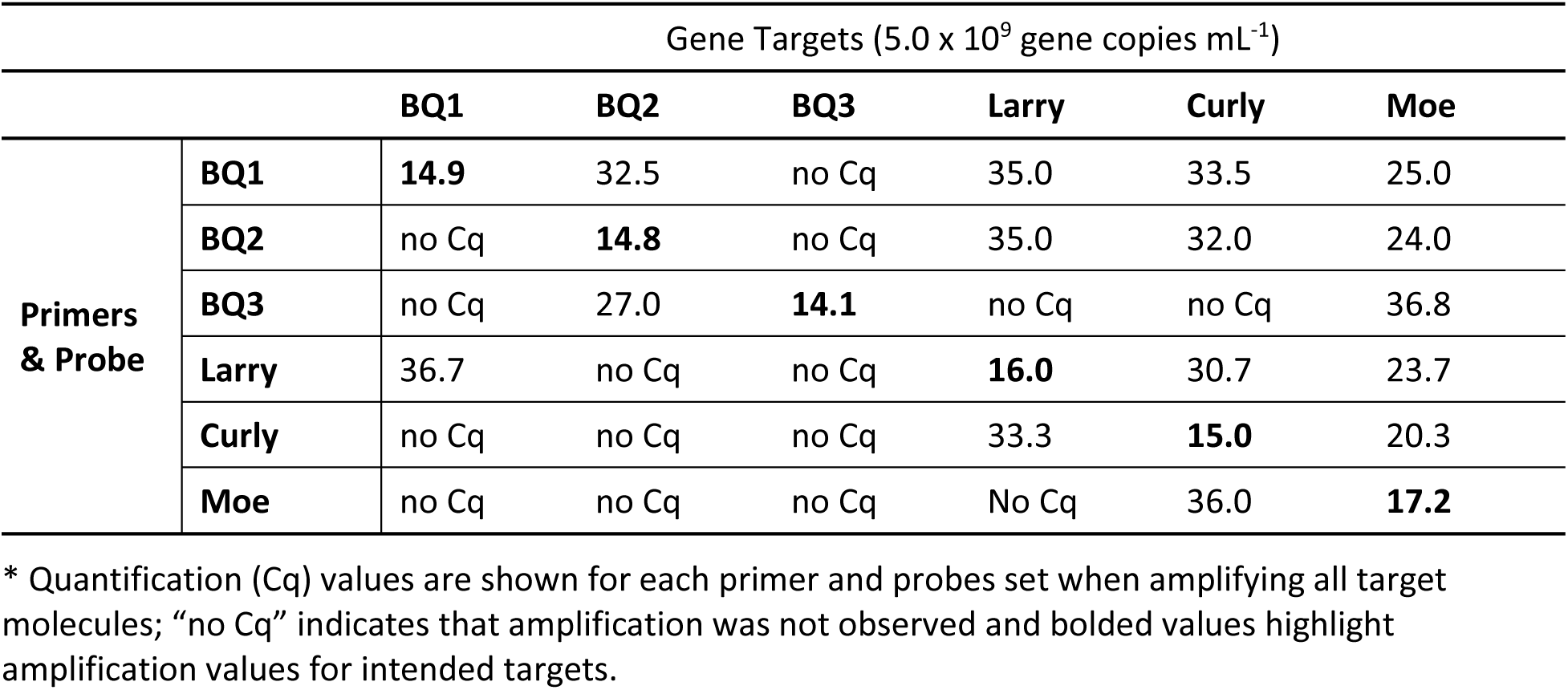
Cross reactivity gene-specific qPCR primers and probes*.

### 3.2 Virus Isolation

To isolate the individual *Chrysochromulina parva* viruses, a virus stock containing a mixture of all known viruses and PLVs (i.e., BQ1, BQ2, Larry, Curly and Moe), as confirmed via qPCR, was propagated through multiple serial infections of *C. parva* culture. We hypothesized that fitness differences between these viruses when grown in specific laboratory conditions would result in the enrichment of certain viruses after several generations of infection. After 5 generations of infection in both parallel propagation series (i.e., 0.45 vs 0.20 μm pore-size filtration), all viruses except for Moe became undetectable by quantitative PCR yet filtered medium from each series lysed *C. parva* cultures.

A portion of the lytic sample collected from the 0.20 μm pore-size filtration was further filtered through a 0.10 μm pore-size filter. The 0.10 μm filtrate contained Moe at high abundance (Cq = 19.5), yet it did not cause culture lysis. On the other hand, when the 0.10 μm filter itself was soaked in sterile DY-V medium overnight, and a 1/10 v/v of this was added into a mid-log phase culture, *C. parva* lysis was observed. None of the known viruses were detected in this culture lysate.

We then used two MPN assays, or ‘dilution to extinction’ approaches, to further purify and isolate the lytic agent present in the fifth-generation lysates from each of the parallel (0.20 versus 0.45 μm filtrations) propagation series. From each MPN assay, a sample at the highest dilution level that produced visible cell lysis was selected for further analysis and propagation. These two samples were propagated through an additional four generations of propagation following the same filtration scheme for the resulting lysates. Of all known viruses, only Moe was detectable in the lysates derived from the 0.20 μm filtration series, whereas the none of the known viruses (i.e., BQ1, BQ2, Larry, Curly, and Moe) were detectable in the 0.45 μm filtration series. Nucleic acids were extracted from a portion of the filtered lysate from 0.45 μm filtration series and were sent for Illumina sequencing.

### 3.3 Sequence Analysis & Phylogeny of an Unknown Lytic Agent

Sequence analysis of nucleic acids recovered from the last generation of lysate from the 0.45 μm filtration series is ongoing. Preliminary analysis resulted in the assembly of four large (i.e., > 50 kbp) contiguous sequences which, via BLAST analysis, were most closely related to the published genome of CpV-BQ2 [52] and represent the genome sequences of a newly discovered *C. parva*-infecting virus which we have labelled CpV-BQ3. Because these contigs cannot be assembled into a single virus genome, and will be further analysed in a future study, they are not reported here. However, several hallmark virus sequences including those encoding the genes PolB (DNA polymerase family B), A32 (packaging ATPase), and VLTF-3 (Poxvirus Late Transcription Factor) were identified in two of the largest contigs. The PolB and VLTF-3 genes were identified within a single 270 kbp contig, whereas the A32 gene was identified within a separate 75 kbp contig. These gene sequences have been submitted to GenBank and have been assigned the accession numbers PX584302 (PolB), PX584303 (A32), and PX584304 (VLTF3). BLAST comparisons of these sequences to sequences in GenBank demonstrated that all three sequences are nearly identical to sequences of the *C. parva*-infecting virus CpV-BQ2, or *Tethysvirus ontarioense*. The DNA sequence of BQ3’s PolB gene is 93 % identical over 3,865 bp to the BQ2 PolB gene, its A32 gene is 95% identical over 839 bp, and its VLTF3 gene is 93 % identical over 1,148 bp.

Phylogenetic analyses of these gene sequences corroborate earlier analyses of members of the *Nucleocytoviricota* like *Tethysvirus ontarioense* (BQ2), reinforcing the monophyly of haptophyte (i.e., prymnesiophyte)-infecting viruses within the *Mesomimiviridae.* The PolB genes of BQ3 and BQ2 were placed as sister taxa in a clade most closely related to *Phaeocystis globosa* virus 14T and *Prymnesium kappa* virus, and the *Chrysochromulina ericinia* virus CeV-01B, or *Tethysvirus raunefjordense* (Figure 1). This same close relationship of BQ3 sequences to genes from other marine haptophyte-infecting viruses is echoed in the phylogenies of the A32 and VLTF-3 genes (Figures S1 and S2, respectively).

**Figure 1.**
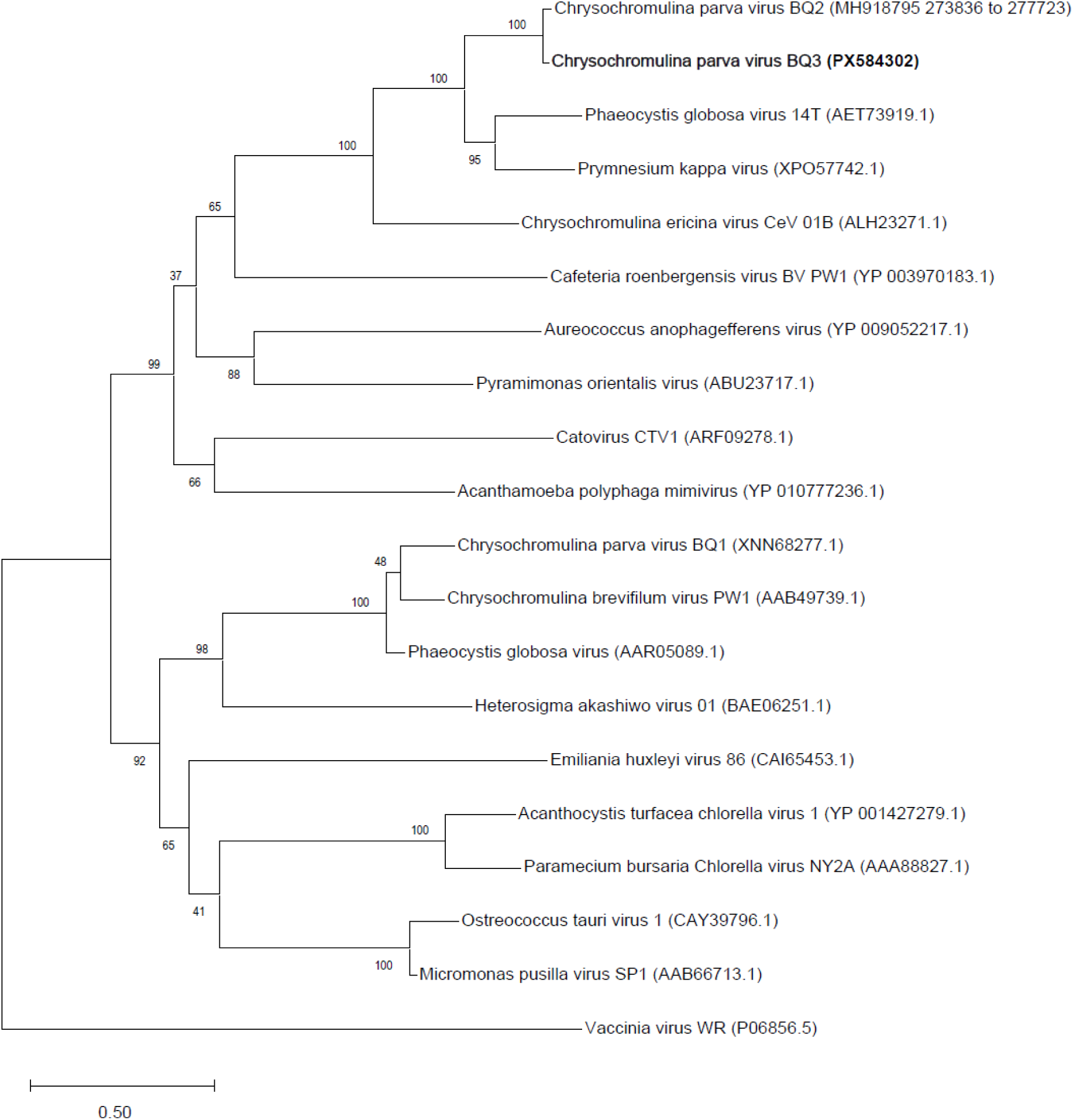
Phylogenetic analysis of virus PolB genes. The phylogeny was inferred using the Maximum Likelihood method and Jones-Taylor-Thornton model of amino acid substitutions [47] and the tree with the highest log likelihood (-44,597) is shown. The final dataset for analysis included 20 amino acid sequences with 2,380 positions.

**Figure 2.**
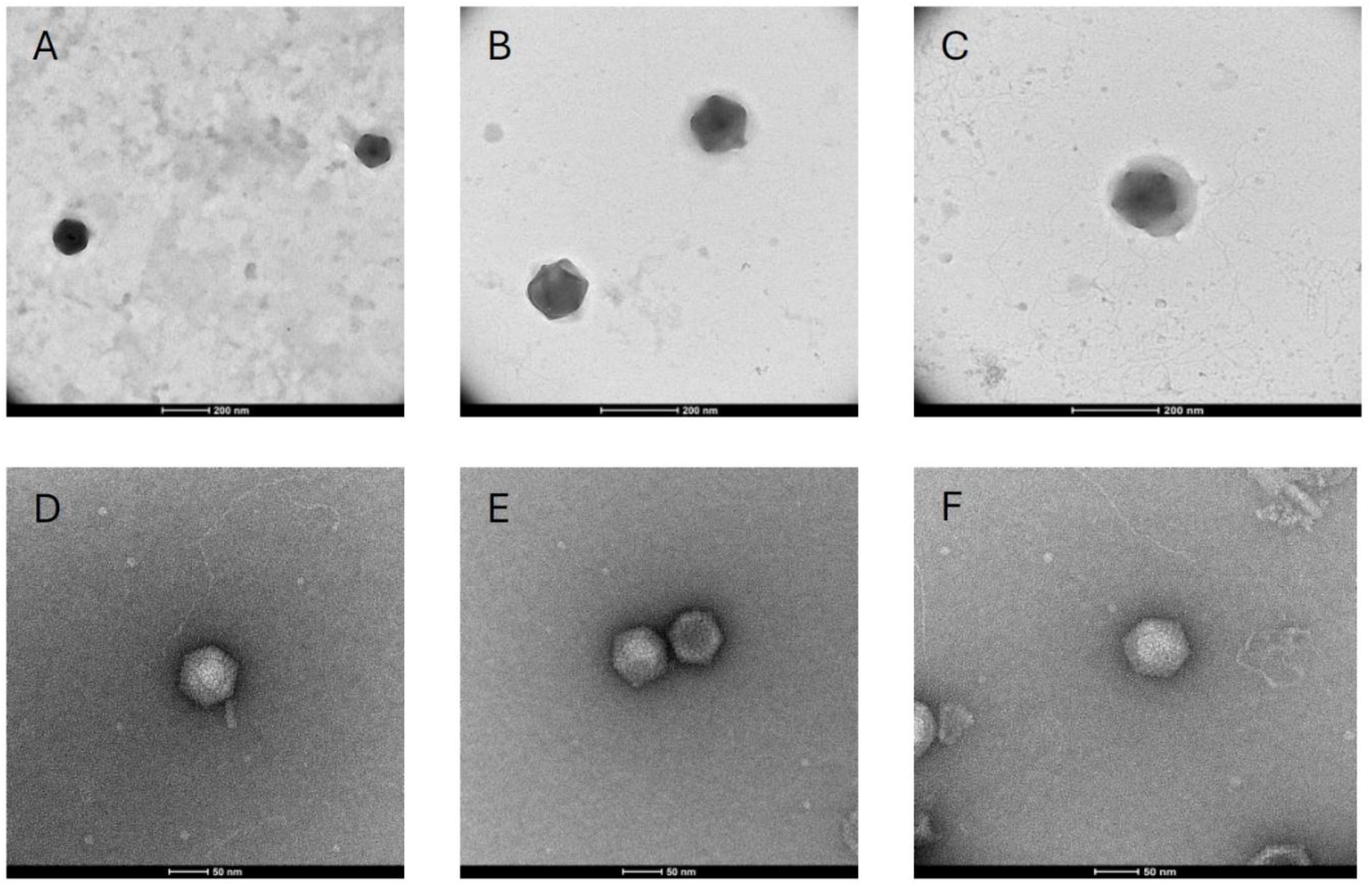
Transmission electron microscopy of viral lysates. (A - C) Micrographs obtained from samples with highly abundant CpV-BQ3 detected with qPCR. (A) Image of ∼150 nm diameter particles in a sample stained with 2 % uranyl acetate on a poly-L-lysine treated grid prepared using the viral lysate that was sent for next-gen sequencing and viewed at 36,000 X magnification. (B-C) Images of ∼150 nm diameter particles observed in samples stained with 0.5% uranyl acetate and viewed at 57,000 X magnification. (D – F) Micrographs obtained from samples with highly abundant CpVV-Moe detected with qPCR stained with 0.5% uranyl acetate and viewed at 120,000 X magnification. Particles range from 60-70 nm diameter depending on the image and axis of measurement.

### 3.4 TEM

Throughout the process of virus isolation, filtered lysate samples were examined using transmission electron microscopy. Examination of the final lysate obtained from the 0.45 μm filtration series, and from which a portion was used for DNA extraction and sequencing, revealed numerous particles with roughly the same icosahedral shape and diameter. The largest particles observed in these grids were approximately 150 nm in diameter (Figure 1A & B) and smallest particles of approximately 140 nm diameter (Figure 1C). Some particles appeared to be surrounded in an amorphous outer layer of material associated with a more rigid appearing core that is similar in size to particles lacking this layer (Figure 1C). Particles with appearances like these ∼150 nm particles (and with both naked and sheathed appearance) were also observed in preparations from earlier stages of isolation, and retrospectively, all these samples contained highly abundant BQ3 PolB genes, as inferred by qPCR. On the other hand, TEM visualization of samples from the 0.20 μm filtration series for which only Moe was detectable by qPCR (prior to the discovery and sequencing of BQ3) were dominated by negatively stained, icosahedral particles approximately 70 nm in diameter across the longest axis (Figure 1 D-F). All these smaller particles appeared to be rigid naked particles, with no outer material associated with them.

### 3.5 Growth experiments

To facilitate visual comparisons of data from all growth experiment trials, the relative abundance of *C. parva* cells were plotted by expressing the abundance of cells in experimental treatments as a proportion of the abundance of cells in the no-infection controls for each trial (Figure 3). In every experimental trial, the infection with BQ3 alone caused complete lysis of *C. parva* cultures within 5 days post infection. However, infections with Moe alone did not appear to cause lysis as cell abundances remained close to the no-infection, growth controls over 20 days (Figure 3A). However, depending on the proportion of Moe in the mixed infection treatments, relative cell abundances differed from the BQ3 infections. Relative cell abundances in the experimental treatments with the highest proportions of Moe (1:1000 and 1:900 BQ3: Moe) remained high throughout the experiment with final abundances above 80% of the abundance of the growth controls (Figure 3B). Cells treated with BQ3: Moe added at 1:500 appeared to experience some lysis with abundances dropping to just over 40% by the fifth day of the experiment and ending at around 20%. Relative cell abundances in treatments with the lowest proportions of Moe (1:10 and 1:200 BQ3: Moe) were similar to infections with BQ3 alone, except cell growth appeared to recover slightly by the end of the experiment with abundances reaching approximately 20% of the growth control on day 20 (Figure 3B).

**Figure 3.**
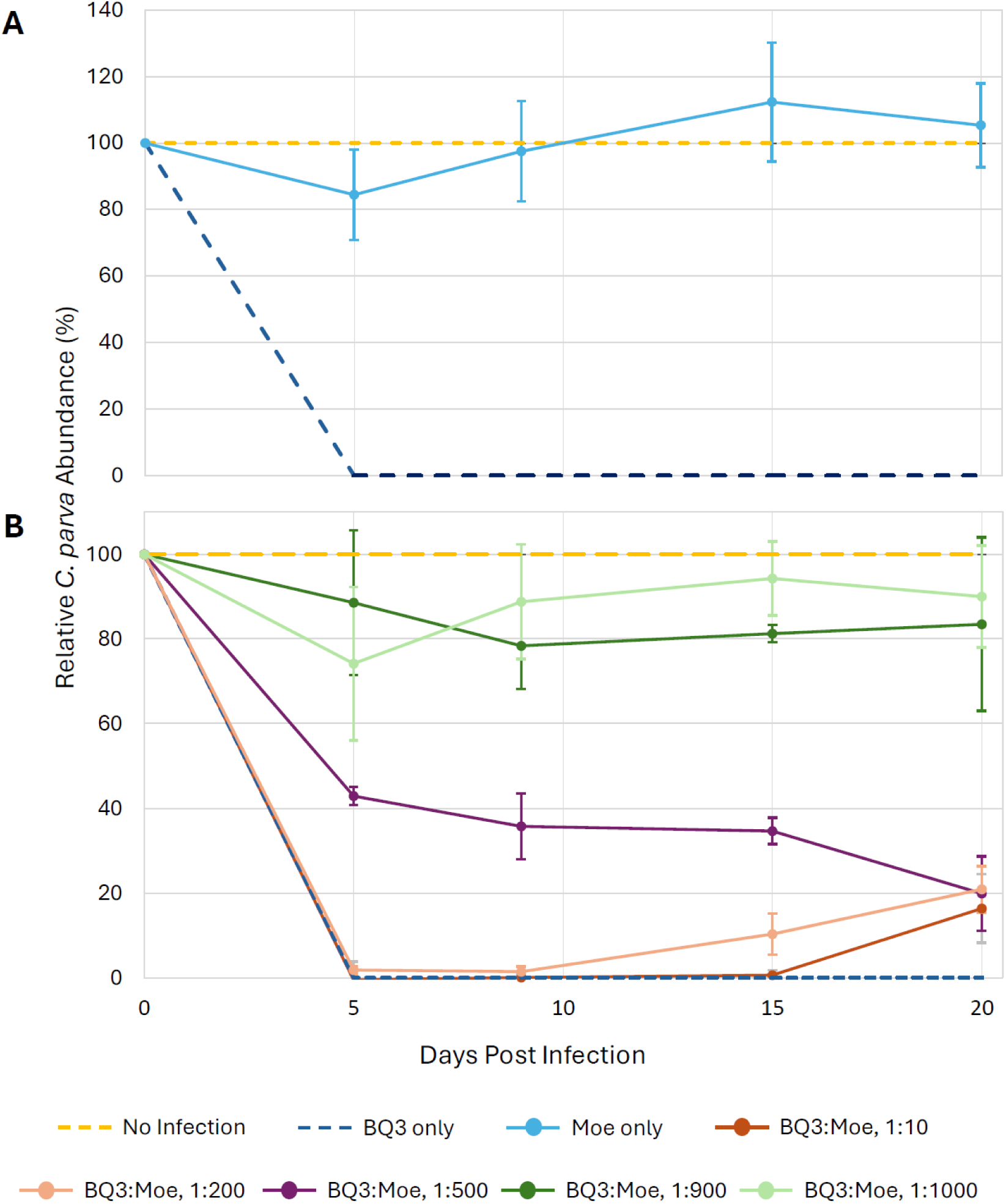
Relative growth of *C. parva* cultures over 20 days. Cell growth was expressed as a percentage of the mean growth observed in triplicate no-infection control flasks; cell counts were normalized across 4 experimental trials utilizing the equation: 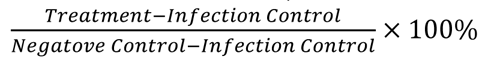. (A) Relative *C. parva* growth when only CpV-PLV Moe was inoculated into triplicate flasks. (B) *C. parva* growth when CpV-PLV Moe was inoculated at different ratios relative to CpV-BQ3 as inferred from qPCR. In both panels, error bars correspond to the propagated standard deviation of growth estimates from triplicate flasks. Standard deviations for each treatment were derived using propagation of uncertainty formulae to account for the standard deviations of *C. parva* abundances observed in replicates of the no-infection controls (dotted yellow lines) and the BQ3-only infections (dotted blue lines); i.e., the error bars shown are larger than they would be if they were derived from treatment replicates alone.

Statistical analyses of cell growth were performed on the raw abundance data (Figure S3) on the 9^th^ day of each experimental trial. Welch’s ANOVA detected significant differences in day-9 cell abundances among treatments within each experimental run, and Games–Howell post-hoc tests revealed clear pairwise patterns. In the first experimental trial which included the BQ3: Moe 1:1000 treatment, cell abundances in the mixed infection were significantly higher than the BQ3 infection (p < 0.001), and cell abundances in replicates with Moe alone were significantly different from the BQ3 infection flasks (p < 0.001). The day 9 cell abundances in the BQ3: Moe 1:1000 and Moe only treatments did not differ significantly from cell abundances in the no-infection growth control replicates (p = 0.14). In the second experimental trial with BQ3: Moe 1:900, and BQ3: Moe 1:10 treatments, day 9 cell abundances in the 1:900 treatment were significantly higher than those of the BQ3 infection control (p < 0.001) but were statistically indistinguishable from the no-infection growth controls (p = 0.13). However, abundances in the BQ3: Moe 1:10 treatment flasks were significantly different from the no-infection control (p < 0.001) but were not statistically different from the corresponding BQ3-alone infections (p = 1.0). In the third experimental trial which included the BQ3: Moe 1:500 mixed infection, cell abundances in the 1:500 flasks were significantly different from both the no-infection growth control (p = 0.026) and the BQ3 infection control (p = 0.001). In the final experimental trial, which included the BQ3: Moe 1:200 mixed infection, cell abundances in the mixed infection differed significantly from the no-infection control (p < 0.001) but did not differ statistically from the corresponding BQ3 infection control (p = 0.23).

In all experimental trials, the abundance of Moe, inferred from qPCR, increased after the initial infection (i.e., day 0) in all treatments which included a mixed infection with Moe and BQ3 (Figure 4). However, after an initial approximately 10-fold increase, the abundance of Moe gene copies decreased back to the original starting abundance in both the 1:900 and 1:100 BQ3: Moe mixed infections. On the hand, Moe gene abundances in the 1:10, 1:200, and 1:500 mixed infections increased by approximately 4.5, 3, and 2 orders of magnitude, respectively, after the 20-day incubation. Finally, the abundance of Moe MCP gene copies decreased from approximately 10^8^ copies mL^-1^ to approximately 1 x 10^7^ copies mL^-1^ in the replicate flasks with infected with Moe alone.

**Figure 4.**
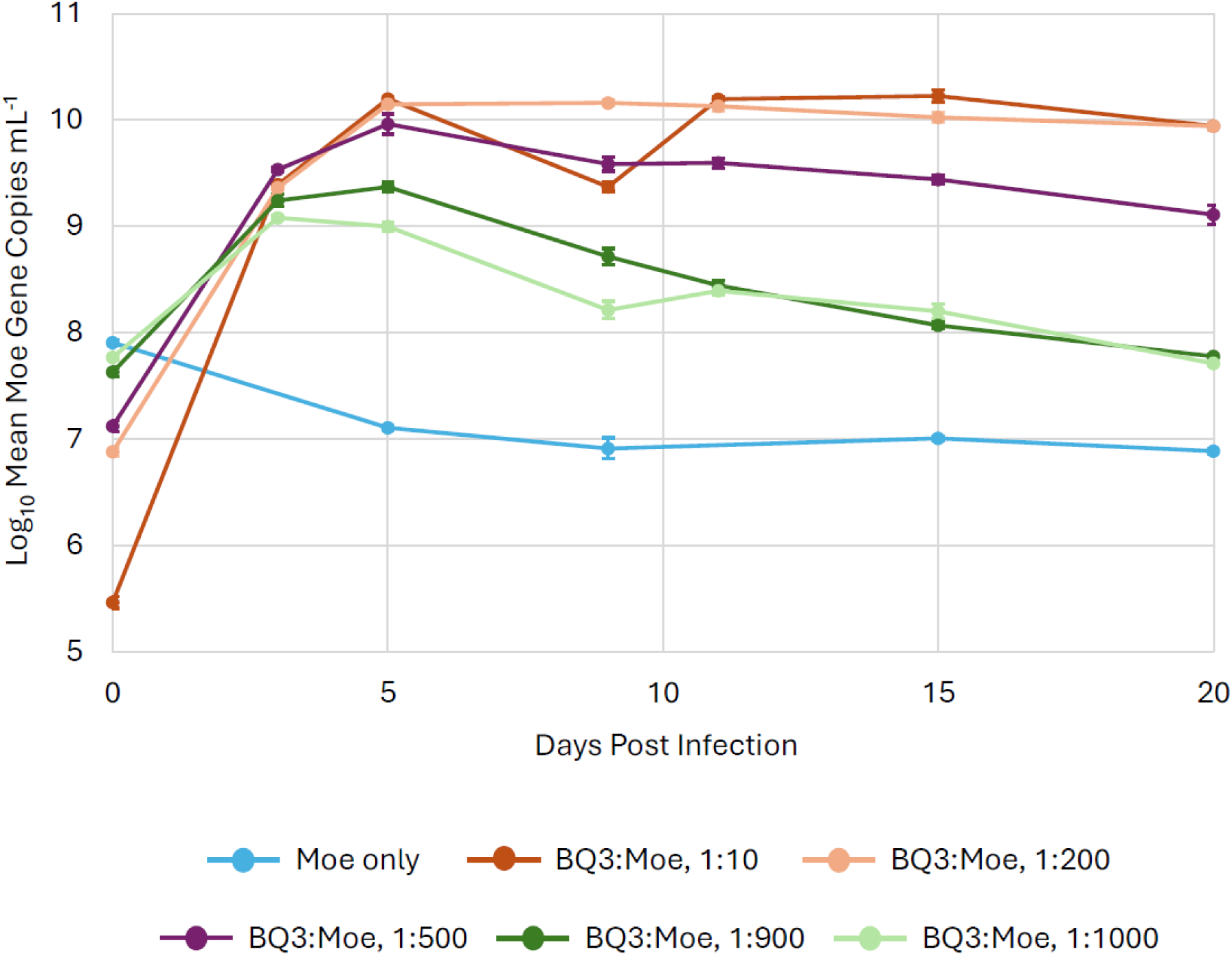
The replication of CpV-PLV Moe in mixed infections. Moe’s replication varied in triplicate experimental flasks inoculated with different ratios of Moe relative to CpV-BQ3 which was held at a constant multiplicity of infection of 0.01 infectious units per cell. The legend shows the approximate ratio of BQ3: Moe for each treatment based on qPCR estimates. The error bars correspond to the standard deviation of Moe gene copies estimated in triplicate flasks, and where they are not visible, they were smaller than the data marker.

Across all trials, the presence of Moe resulted in significantly reduced BQ3 replication on Day 9; i.e., BQ3-only infections resulted in significantly higher BQ3 gene copy abundances than the corresponding mixed-infection treatments. Specifically, BQ3 abundances in the BQ3-only infections were significantly higher than BQ3: Moe 1:1000 (p < 0.001), BQ3: Moe 1:900 (p < 0.001), BQ3: Moe 1:500 (p < 0.001), BQ3: Moe 1:200 (p = 0.0012), and BQ3: Moe 1:10 (p < 0.001).

To help visualize the effect of Moe on the replication of BQ3, the fold-change reduction of BQ3 gene copies in the mixed infections relative to the infection with BQ3 alone was plotted (Figure 5). In the treatment flasks with the highest proportion of Moe relative to BQ3 (BQ3: Moe 1:1000 and 1:900 mixtures) the replication of BQ3 was reduced approximately 300-fold [antilog_10_(2.5) = 316], whereas lower proportions of Moe resulted in lesser impact on BQ3 replication. BQ3 replication was reduced approximately 50-fold in the BQ3: Moe 1:500 flasks, 3-fold in the 1:200 mixture, and about 2-fold in the 1:10 mixture of BQ3: Moe.

**Figure 5.**
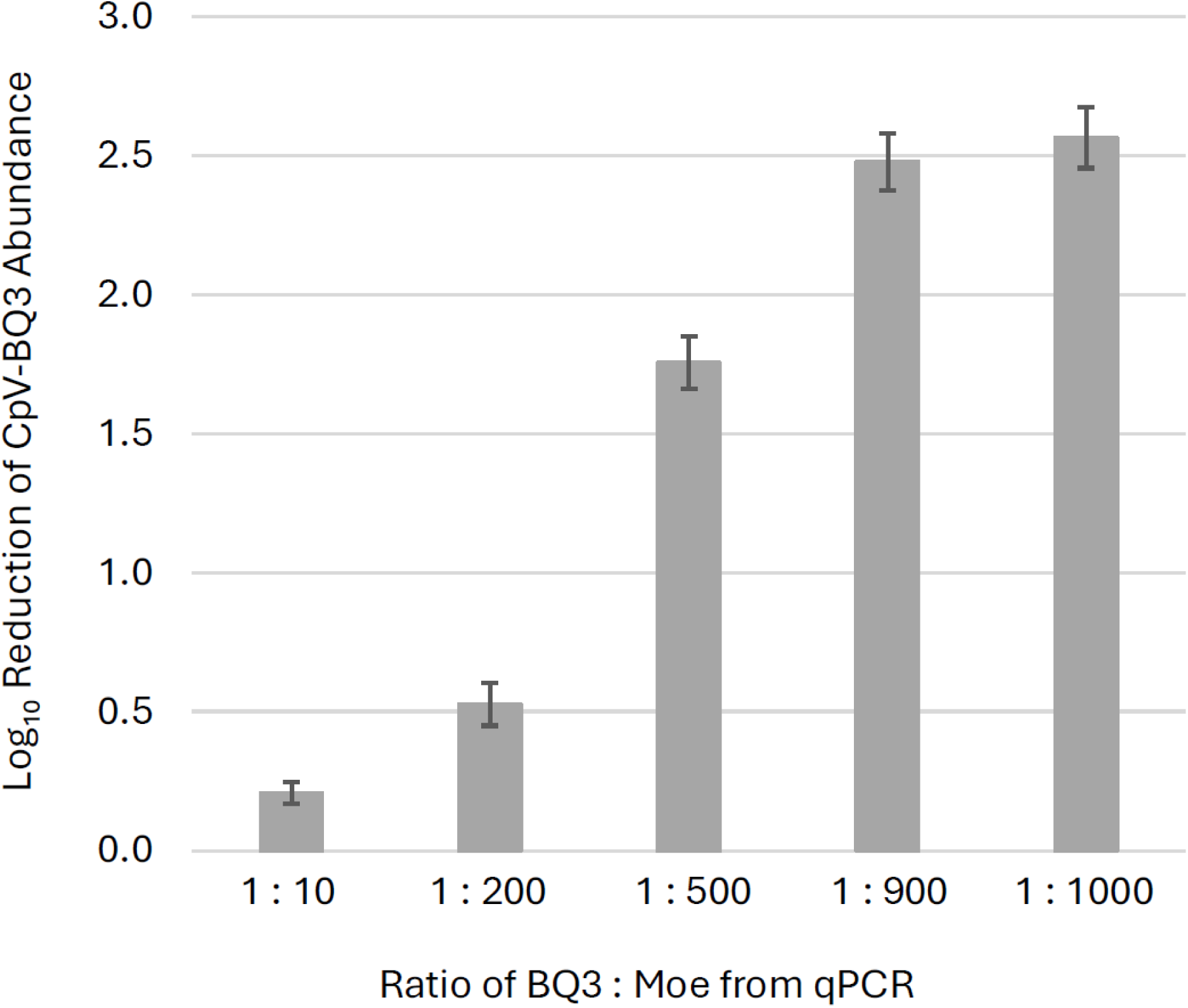
Inhibition of CpV-BQ3 replication. The fold-change reductions of BQ3 abundances in mixed infections relative to corresponding BQ3-only infections are shown for different ratios of CpV-BQ3: CpV-PLV Moe after 9-days post inoculation. Estimates represent the mean fold-reduction of BQ3 abundance in triplicate flasks relative to the mean in triplicate BQ3-infection controls. The error bars correspond to the standard deviation of the estimates, and ratios of BQ3: Moe were based on qPCR estimates of virus abundances.

Welch’s ANOVA demonstrated a significant difference in BQ3 reduction across treatments. A Games–Howell post-hoc test demonstrated that the only non-significant comparison was between the 1:900 and 1:1000 BQ3: Moe treatments (p = 0.637), indicating that these two mixtures resulted in similar levels of BQ3 reduction. All other pairwise comparisons resulted in statistically significant differences in the reduction of BQ3 replication. Relative to the 1:10 treatment, the replication of BQ3 was significantly reduced in the 1:1000, 1:200, 1:500, and 1:900 treatments (p < 0.001 in all cases). Additional pairwise comparisons revealed significant differences among the treatments with intermediate ratios of the BQ3: Moe mixed infections, including 1:1000 vs. 1:200 (p < 0.001), 1:1000 vs. 1:500 (p < 0.001), 1:200 vs. 1:500 (p < 0.001), 1:200 vs 1:900 (p < 0.001), and 1:500 vs 1:900 (p = 0.002).

To determine if the effect of Moe on the replication of the putative Phycodnavirus CpV-BQ1 was similar to its effects on the Mesomimivirus CpV-BQ3, an additional coinfection experiment was conducted with a 1:500 mixture of BQ1 and Moe. This coinfection resulted in a different lysis pattern compared to the infections with a 1:500 mixture of BQ3: Moe (Figures 6A and 3B). In the mixture with BQ1, *C. parva* abundances remained close to 80% of the no-infection control until the 20^th^ day of the experiment when cell abundances dropped to approximately 30 ± 4% of the *C. parva* abundances in the no-infection control.

**Figure 6.**
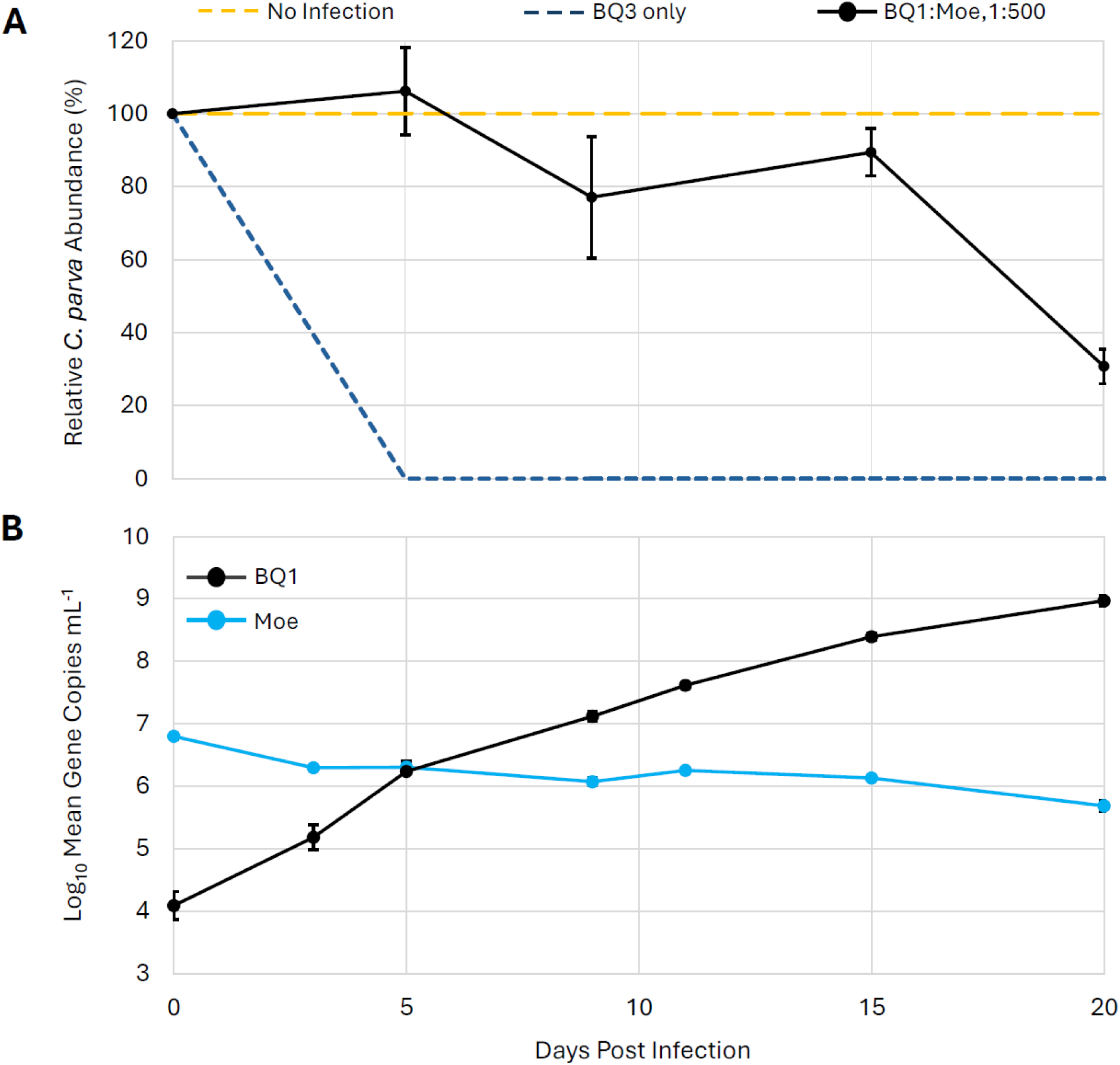
Replication of CpV-PLV Moe in the presence of the virus CpV-BQ1. (A) C. parva growth is shown as a percentage relative to no-infection control flasks. Inoculation with CpV-BQ3 alone was used an infection control. (B) The abundances of CpV-BQ1 and CpV-PLV Moe were estimated using qPCR and are shown for triplicate flasks inoculated with both BQ1 and Moe at a ratio of 1: 500, BQ1: Moe. The error bars represent standard deviation of triplicate flasks, and where they are not visible, they were smaller than the data marker.

The abundances of CpV-BQ1 and CpVV-Moe in the BQ1: Moe mixed infection were monitored over the 20-day sampling period by examining the abundance of viral gene copies via qPCR (Figure 6B). CpV-BQ1 appeared to replicate throughout the sampling period of the experiment, steadily increasing from the initial inoculum of 1.3 × 10⁴ gene copies mL⁻¹ up to 9.5 × 10⁸ gene copies mL⁻¹ on day 20. In contrast, there was no evidence of Moe’s replication as its abundance decreased slightly from an initial inoculum of 6.4 × 10⁶ gene copies mL⁻¹ to a final abundance of approximately 5.0 × 10⁵ gene copies mL⁻¹.

## 4. Discussion

### 4.1 Validation of qPCR methods

The motivation for this study was to determine the relationship between the algal viruses CpV-BQ1 and BQ2 and the polinton-like viruses CpV-PLV Larry, Curly, and Moe. To accomplish this, these viruses needed to be purified from the mixed cultures in which they had been propagated in since their discovery; all originated from a single Lake Ontario, Canada water sample collected in 2011 [37]. Efforts to isolate individual viruses were facilitated by gene-specific qPCR which allowed the abundance of each virus to be monitored independently, even in mixed infections. Quantitative PCR assay validation experiments demonstrated that all primer and probe sets performed with appropriate specificity when amplifying cloned gene fragments from purified plasmid preparations. To provide a conservative estimate of the signal generated from off-target amplifications, the concentration of plasmid templates used in validation experiments was generally higher than observed in experimental infections; only Moe MCP genes exceeded 10^9^ gene copies mL^-1^ in any experimental infection (Figure S3). The least specific primer and probe set (i.e., generating the most off-target amplification) targeted Curly MCP. This set amplified Moe MCP genes, albeit with approximately 40-fold reduced sensitivity based on the Cq difference of 5.3 between off-target and on-target amplification. Most importantly for the experimental infections described here which focused on interactions of BQ3 and BQ1 with Moe, the primers and probes for these templates were all at least 1000 times less sensitive to non-target templates (i.e., Cq difference ∼ 10), and the primers and probes for Moe did not amplify any off-target genes except Curly and that with nearly a million-fold reduced sensitivity (Cq difference was ∼19).

Quantitative PCR of known viruses (BQ1 and 2, and Larrry, Curly, and Moe) were used to monitor changes in virus abundance through serial propagation and dilution-to-extinction experiments and provided evidence that virus stocks included a previously unknown virus. In the lineage propagated after filtration through 0.45 µm pore-size membranes, all known viruses dropped below detection. On the other hand, the lineage filtered through 0.2 µm membranes, only Moe remained detectable, maintaining relatively high abundances in the lysates (∼1.0 x 10^9^ copies mL^-1^). Together, these results stimulated the hypotheses that Moe was able to replicate on its own, or viral lysates contained another lytic agent that hadn’t been discovered in earlier characterizations of *C. parva* viruses [36, 37]. To test these hypotheses, we sequenced nucleic acids from a lysate with no detectable viruses and conducted the mixed-infection experiments described herein.

### 4.2 Discovery of a new isolate of Tethysvirus

Although analysis of Illumina sequences obtained from the infection series propagated following 0.45 µm filtration and subsequent end-point dilution is still underway, three hallmark viral sequences have been identified in contigs of the newly isolated *C. parva*-infecting virus reported here. For simplicity and consistency with previous *C. parva* isolates from Lake Ontario, Canada, we have informally named this virus CpV-BQ3 because it is the third *C. parva* virus isolated from the Bay of Quinte water sample that led to the isolation and identification of BQ1 and BQ2 [36, 37]. At the time of the current study, the virus CpV-BQ2 has not yet been isolated but remains detectable in some mixed cultures of CpVs. The fact that BQ3 seems to outcompete BQ2 in serial propagation experiments suggests it has a fitness advantage over BQ2 when replicating in laboratory conditions. Efforts to purify BQ2 via end-point dilution have not been exhaustive and continued efforts could lead to its eventual purification. Having specific qPCR assays for each virus will facilitate these efforts allowing work to focus on lysates enriched in BQ2.

Maximum likelihood phylogenies of the PolB, VLTF3, and A32 genes of BQ3 (Figure 1, Supplementary Figures 1 and 2) demonstrated the close relationship of all three genes with genes of *Tethysvirus ontarioense*, or CpV-BQ2. Considering the criterion that members of the same species have pairwise average nucleotide identities (ANI) above 95% for more than 75% of the predicted genes in each genome [32], it remains unclear if CpV should be considered a strain of *T. ontarioense*, or if it should be considered a new species of *Tethysvirus*. Nevertheless, these phylogenies unambiguously placed genes of BQ3 as sister to BQ2 genes, and moreover, that the relationships between BQ2 and BQ3 and other haptophyte-infecting mesomimiviruses of *Phaeocystis globosa* and *Haptolina ericinia* (syn. *Chrysochromulina ericinia*) were consistent with established phylogenies [e.g., 21, 32, 33, 37, 53]. Hence, it is tenable to propose that CpV-BQ3 is a species of *Tethysvirus*, a *Varidnaviria* genus in the family *Mesomimiviridae*, order *Imitervirales*, class *Megaviricetes*, kingdom *Bamfordvirae.* Using the PolB gene sequence of BQ3, a new qPCR primer and probe set was designed and validated, providing a tool to monitor its abundance in lysates generated from mixed virus infections.

### 4.3 Virion imaging via TEM

A portion of the final lysate derived from serial propagation of 0.45 µm filtered lysates, and from which nucleic acids were extracted for Illumina sequencing, was imaged using transmission electron microscopy. In these samples, a single particle morphology was observed corresponding to an approximately 150 nm diameter icosahedral particle. Interestingly, some, but not all, particles observed in the same grids appeared to be surrounded by an outer sheath layer (Figure 2B & C). Additionally, as noted in the results, retrospective examination of samples using the newly designed BQ3 qPCR primers and probes demonstrated that earlier samples from this lineage also contained similar sized particles with the same morphologies, i.e., both naked and sheathed particles were observed in different TEM micrographs of lysates with highly abundant BQ3 (Supplementary Figure 4). It is not clear whether the ‘sheath layer’ is a structural component of the virions, like the fiber layer observed in structural studies of Mimivirus [54], but the fact that this layer has been seen in different preparations of BQ3 provides evidence that it is a virion component rather than a artefact. Further experimental infections and characterization using cryo-EM visualization purified BQ3 virions will be needed to resolve the structure of this unusual outer layer and determine if it is an essential component of virions required for infection.

An additional series of lysates were propagated after filtration through 0.2 µm pore-size filters. Quantitative PCR analysis of these lysates demonstrated that the only known virus present was the polinton-like virus CpV-PLV Moe. Imaging this sample with TEM revealed the presence of virions with icosahedral morphology approximately 70 nm in diameter. These particles were more easily negatively stained than BQ3 particles, and appeared as well-defined, naked particles similar in morphology and size to other *Preplasmiviricota* [17] virions like PLV Gezel-14T which co-parasitizes the *Phaeocystis globosa* along with the *Mesomimiviridae* PgV-14T [34], the Mimivirus virophages Sputnik and Zamilon, and the CroV virophage Mavirus [11]. Earlier samples from this lineage also revealed the presence of particles corresponding to both BQ3 and Moe (Supplementary Figure 4). When lysate samples with highly abundant Moe gene fragments were passed through 0.1 µm pore-size filter and the resulting filtrate was added to mid-log phase culture of *C. parva*, no lysis was observed yet Moe remained highly abundant. On the other hand, material retained on this 0.1 µm filter was lytic and infectious providing evidence for the existence of an unknown virus, and one that presumably supported replication of the putative polinton-like virus, CpV-PLV Moe. Following sequencing of BQ3 and development of a BQ3-specific qPCR assay, we returned to samples from the lineage that led to the TEM images of Moe shown here and were able to amplify BQ3 PolB genes, albeit much lower abundances than Moe. Thus, it seems that filtrations through 0.2 µm pore-size filters enriched lysates for Moe but also permitted passage of the ∼150 nm BQ3 particles.

### 4.4 Mixed Infections Reveal the Hyperparasitic Replication of Moe

Learning that the 0.1 µm pore-size filtration eliminated lytic activity but did not eliminate qPCR detection of Moe MCP genes, stimulated the mixed infection experiments to determine how Moe replicates. Before discussing the results of the infection experiments reported herein, it important to consider the statistical approaches used in the analyses and described in the results. Logistical constraints of this study were related to the space available in growth chambers and our ability to collect samples at each timepoint from replicate flasks of each treatment while minimizing the sampling impact on multi-day incubations. These constraints required treatments to be conducted in separate experimental trials, each accompanied by positive and negative infection controls to permit comparisons across trials. Thus, treatment replication was limited to triplicate incubations for treatment. Since treatment replication was limited to triplicates, formal tests of normality could not be performed reliably; therefore, observed viral and cell abundances were assumed to approximate normality. Within each experimental trial, all replicates were inoculated using the same viral stock, ensuring consistency of initial conditions and supporting this assumption despite the small sample size. For analyses employing ANOVA comparisons, residuals were normalized using power transformations (i.e., square-root or fractional power transformations), with appropriate transformation selected to satisfy model assumptions based on model diagnostics (Q–Q plots). Variance homogeneity was not assumed for any comparisons as mixed infections with higher ratios of Moe to BQ3 resulted in larger variances in cell and virus abundances. Accordingly, Welch’s tests were used instead of equal-variance alternatives providing a more conservative and robust test for these comparisons. When evaluating the effect of the virophage Moe on BQ3 replication, a Bonferroni-adjusted significance threshold (α = 0.025) was applied to account for multiple comparisons; each experimental trial involved either one or two comparisons between the BQ3 infection control and the corresponding co-infection treatment.

Overall, mixed infections demonstrated that Moe depended on its *Mesomimividae* host - BQ3. When Moe was added to *C. parva* cultures alone, the cultures continue to grow at abundances that did not differ from mock-infected control cultures (Figure 3A, Supplementary Figure 3A). Using qPCR to track Moe abundances in these infections demonstrated that Moe did not replicate. Indeed, when *C. parva* was infected with Moe alone, Moe’s abundance dropped 10-fold over first 5 days of the incubation and remained at this reduced abundance throughout the course of the 20-day incubation (Figure 4). The observed loss of Moe gene copies through this experiment cannot be explained at this time, but it is plausible that Moe particles and amplifiable gene fragments were destroyed during the incubation by various biological and/or abiotic sources of decay that are known to eliminate viruses in natural environments [55]. When Moe was inoculated into *C. parva* cultures alongside CpV-BQ1, a putative Phycodnavirus [35], Moe abundances also dropped approximately 10-fold (Figure 4) whereas BQ1 gene abundances increased by a factor of approximately 10^5^ (Figure 6B) and replicate *C. parva* cultures lysed, albeit more slowly than cultures inoculated with the putative Mesomimivirus, CpV-BQ3 (Figure 6A).

On the other hand, when Moe was added to *C. parva* cultures in the presence of BQ3, Moe abundances always increased. As noted in the methods section, all mixed infection experiments were conducted at a constant MOI of 0.01 infectious units of BQ3 or BQ1 per cell (estimated from MPN assays and cell counts), but with differing proportions of Moe relative to these viruses. Based on results from preliminary experiments not described here, we hypothesized that Moe affected both *C. parva* growth and virus replication in a density-dependent manner. Hence, we conducted a series of experimental trials which included mixed infections with BQ3 and Moe ranging from 1:10 to 1:1000 gene copies of BQ3: Moe. These proportions were created by varying the inoculum of Moe and were empirically confirmed in all replicate incubations at time zero. In cultures receiving the lowest proportion of Moe (1:10), Moe abundances increased 50,000-fold over the 20-day incubation, whereas in cultures receiving the highest proportions of Moe (1:1000 and 1:900), Moe increased approximately 10-fold during the first five days of the experiment and then dropped back to starting abundances. Incubations with an intermediate proportion of BQ3: Moe (1:500) resulted in an intermediate increase in Moe’s abundance starting at approximately 10^7^ gene copies mL^-1^ and increasing 100-fold to a final abundance of 10^9^ gene copies mL^-1^ by the end of the experiment. Together these results demonstrated that Moe production saturated when incubations were started with high levels of Moe, and that some decay or loss of Moe influenced its final abundances in the 20-day batch cultures. Most importantly though, these experiments provide clear evidence that Moe replicates in the presence of the putative Mesomimivirus CpV-BQ3, but not in the presence of the putative Phycodnavirus CpV-BQ1. This observation is consistent with knowledge of other isolated virophages and PLVs which all replicate with a Nucleocytoviricote helper virus in the order *Imitervirales* [11, 19], and while there is some evidence of virophages associated with phycodnaviruses [6], these associations were based on metagenomics and have not yet been confirmed through cultivation.

Confirming the hyperparasitic replication strategy of Moe raised questions about its effects on the replication of its helper virus and the growth of its host cells. The mixed-infection experiments demonstrated that Moe inhibited BQ3 replication in a dose-dependent manner. Every experimental trial included infections with BQ3 alone and all these single-infection incubations resulted in the rapid lysis of host cells (Figure 3, Supplementary Figure 3) even at the low MOI used for all trials (0.01 infectious units per cell). In mixed infections, BQ3 replication was observed, however, the extent of replication was modulated by the relative abundance of Moe. At lower Moe-to-BQ3 ratios, Moe exerted a comparatively small but statistically significant inhibitory effect on BQ3 replication, and as the relative abundance of Moe was increased to 500 Moe gene copies per BQ3 gene copy, BQ3 replication was notably suppressed, and day-9 BQ3 abundances were approximately 300-fold lower in the 1: 1000 BQ3: Moe mixed infection compared to the corresponding infections with BQ3 alone (Figure 5). Observations of reduced BQ3 replication in mixed infections with Moe were also associated with increased survival of *C. parva*, with some cultures maintaining relatively high cell abundances over the course of the 20-day incubations. For example, in the mixed infections with BQ3: Moe inoculated at 1:1000 and 1:900 proportions, *C. parva* abundances dropped slightly but remaining above 80 % of the abundances observed in the corresponding no-infection control cultures. At lower proportions of Moe in mixed infections (i.e., at 1:10 and 1:200 BQ3: Moe), this protection of *C. parva* growth was less apparent with lysis patterns resembling the BQ3 only infections. Interestingly, C. parva seemed to recover from lysis at these lower Moe mixed infections with cell abundances increasing to 20% of the abundance in the control flasks by the end of the experiments (Figure 3B). Some recovery of *C. parva* was even observed in the BQ3-only infections suggesting the existence of a small population of cells resistant to BQ3 infections (Supplementary Figure 3). Because the experiments described here were based on batch cultures, cell abundances did not continue to rise after this time-point and all cells disappeared within 28 days of the initial inoculum, even for control cultures that were not infected, presumably because of the depletion of nutrients in the growth medium. Nonetheless, this pattern of growth in some infections suggests that infected *C. parva* cultures may be able to recover via a resistance mechanism that has not yet been studied. This phenomenon will be interesting to follow-up in future work on this system.

Collectively, these results indicate a negative relationship between Moe dosage and BQ3 replication, and a positive relationship with *C. parva* growth and survival. Relating the proportion of Moe added in mixed infections to BQ3’s replication suggests that Moe-mediated inhibition of BQ3 replication may follow a sigmoidal relationship, with substantial host protection emerging above some threshold ratio of BQ3: Moe and diminishing further increases in host protection or inhibition of virus replication at higher relative doses of Moe (Figure 5). Consistent with this interpretation, reductions in BQ3 replication did not differ appreciably between Moe-to-BQ3 ratios of 900:1 and 1000:1, or 1:10 and 1:200, and was intermediate at the intermediate ratio of 500:1. However, threshold values for the effect of Moe on cell growth or helper virus replication cannot be determined with this data set as additional mixed infection experiments at intermediate or even higher doses of Moe were not conducted.

## 5. Conclusions

Through our efforts to purify individual viruses within the *C. parva*-virus system, we have discovered a new virus, named CpV-BQ3, that supports the replication of CpV-PLV Moe. TEM imaging of purified Moe particles revealed its morphological similarity to other virophages like Sputnik and mavirus [19] and PLVs such as Gezel-14T [34]. Sequencing and TEM imaging of BQ3 provided evidence that this newly discovered virus should be classified as species of *Tethysvirus*, a genus of the *Mesomiviridae* in the order *Imitervirales*, phylum *Nucleocytoviricota*. Based on phylogenetic analysis of three key virus genes encoding PolB, VLTF3, and A32 proteins, CpV-BQ3 is a close relative of CpV-BQ2, but it remains unclear if it should be considered a strain of *Tethysvirus ontarioense* or a new species. Mixed infection experiments demonstrated that the putative Mesomimivirus BQ3 supports the replication of Moe, whereas the putative Phycodnavirus CpV-BQ1 does not. These experiments also demonstrated a dose effect of Moe where the proportion of Moe relative to its helper virus BQ3 influenced both BQ3’s replication and *C. parva’s* growth. At high ratios of Moe relative to BQ3, BQ3’s replication was inhibited and cell lysis was relieved, allowing *C. parva* cultures to continue growing. This result was somewhat surprising considering experiments Moe’s closest cultivated relative, Gezel-14T, did not demonstrate its ability to rescue its host cellular hosts from lysis [34]. On the other hand, the fact that some virophages like Zamilon do not provide host protection while others like Mavirus and Sputnik do [17], highlights the diverse replication strategies and impact of hyperparasitism among members of the *Polisuviricotina*, the subphylum of the *Preplasmiviricota* that includes virophages and polinton-like viruses [17].

The precise mechanism through which Moe inhibits BQ3 replication remains unclear. Moe may interfere with one or multiple stages of the giant virus replication cycle. Studies of other virophage systems have noted disruption of giant virus morphogenesis resulting in the production of defective particles, or a general inhibition of giant virus particle production by hijacking replication machinery or delaying the appearance of virus factories [11]. Additional studies will be required to determine the mechanisms of Moe’s influences on the replication of its helper viruses and growth of its host cells. It also worth noting here that there is currently no evidence that Moe can exist as a provirophage element in its helper viruses or host cells, as observed for other virophage systems [e.g., 56, 57, 58] including Moe’s close relative, the PLV Gezel-14T [34]. Throughout the experiments described here, and in other work not described, qPCR of purified *C. parva* DNA or BQ3 lysate DNA with Moe primers and probes has never produced detectable amplification. Like the diverse replication strategies of virophages and related satellite viruses, the ability of virophages to integrate into host genomes appears to not be an evolutionarily conserved aspect of these hyperparasitic viruses. Nonetheless, tripartite infection systems like *C. parva*-BQ3-Moe highlight the fascinating and surprising biology of cellular parasites. The fact that CpV-PLV Moe appears to have a density-dependent influence on its helper virus’ replication and host cell survival underscores the complex ecology of these systems and raises even more questions about algal virus evolution and persistence, and the diversity and ecology of viral hyperparasites.

## Author Contributions

Conceptualization, S.M.S. and I.I.; Methodology, S.M.S, I.I., G.T., C.N.P., G.B.; Validation, S.M.S., I.I. C.N.P; Formal Analysis, C.N.P. and G.T.; Resources, S.M.S.; Data Curation, S.M.S., I.I., and C.N.P.; Writing – Original Draft Preparation, S.M.S, G.T., I.I.; Writing – Review & Editing, all authors; Supervision, S.M.S.; Project Administration, S.M.S.; Funding Acquisition, S.M.S. and G.T.

## Acknowledgments

The authors would also like to acknowledge Dr. Lindsey Fiddes, a Microscopy Technician at the Microscopy Imaging Lab of the Temerty Faculty of Medicine at the University of Toronto, for her assistance with transmission electron microscopy. We would also like to acknowledge Ms. Daphne Arguelles for her contributions to the validation studies of the qPCR assays used for this work.

---

**Supplementary Figure 1.**
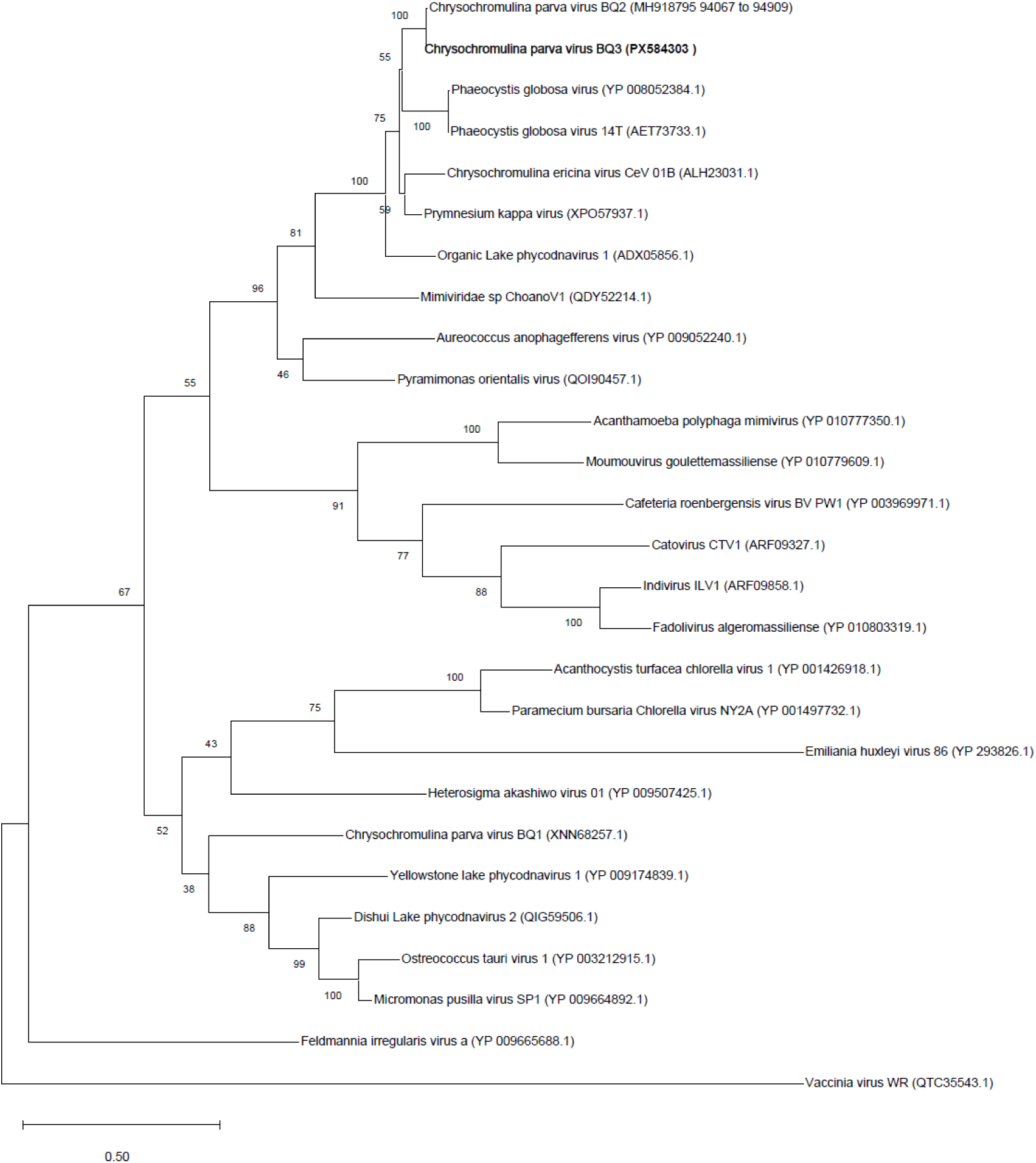
Maximum Likelihood analysis of virus A32 genes. The phylogeny was inferred using Maximum Likelihood and Jones-Taylor-Thornton model [47] of amino acid substitutions and the tree with the highest log likelihood (-10,757) is shown. The final dataset for the analytical procedure included 27 amino acid sequences with 345 positions

**Supplementary Figure 2.**
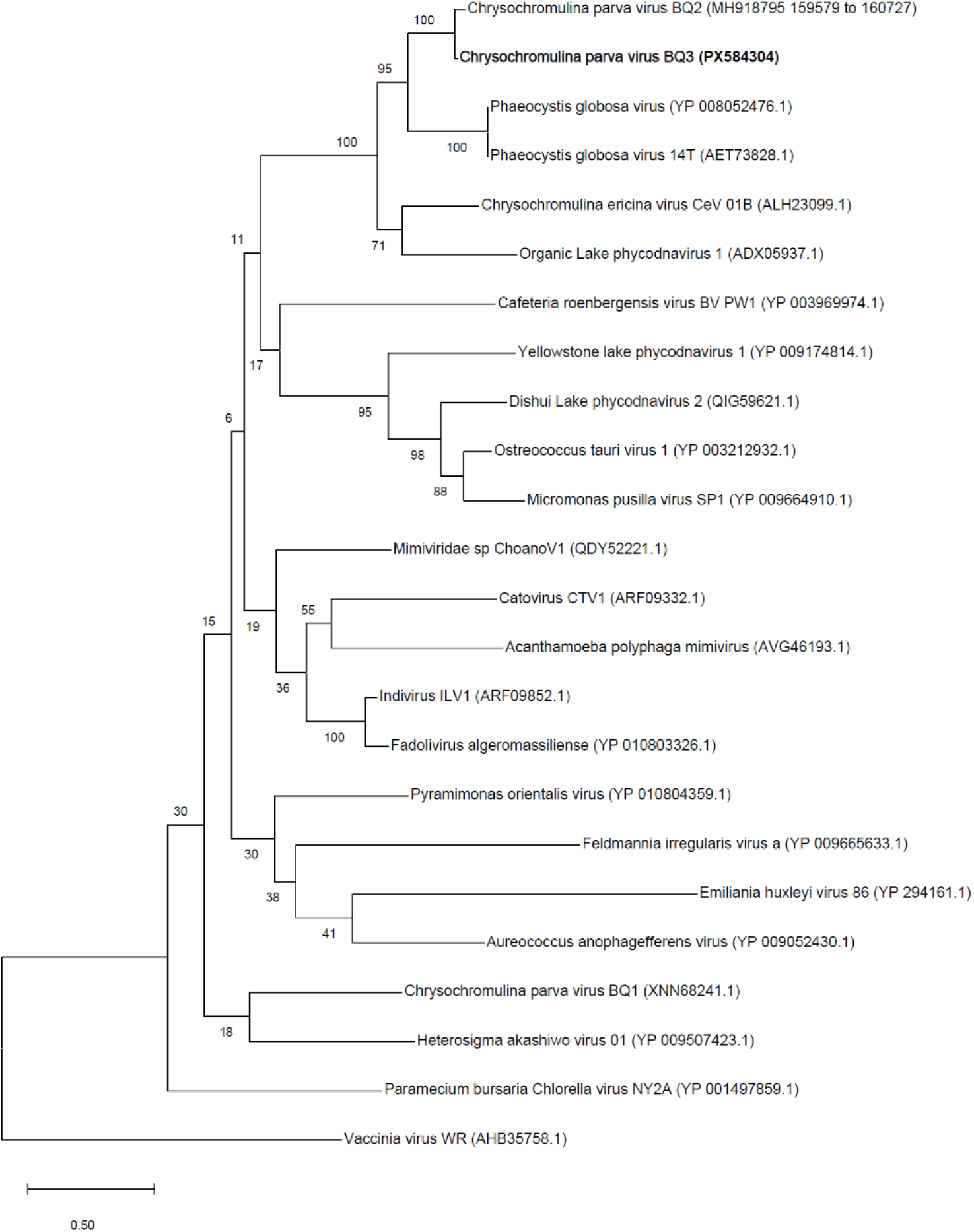
Maximum Likelihood analysis of virus VLTF-3 genes. The phylogeny was inferred using Maximum Likelihood and Jones-Taylor-Thornton model [47] of amino acid substitutions and the tree with the highest log likelihood (-15,024) is shown. The final dataset for the analytical procedure included 24 amino acid sequences with 509 positions.

**Supplementary Figure 3.**
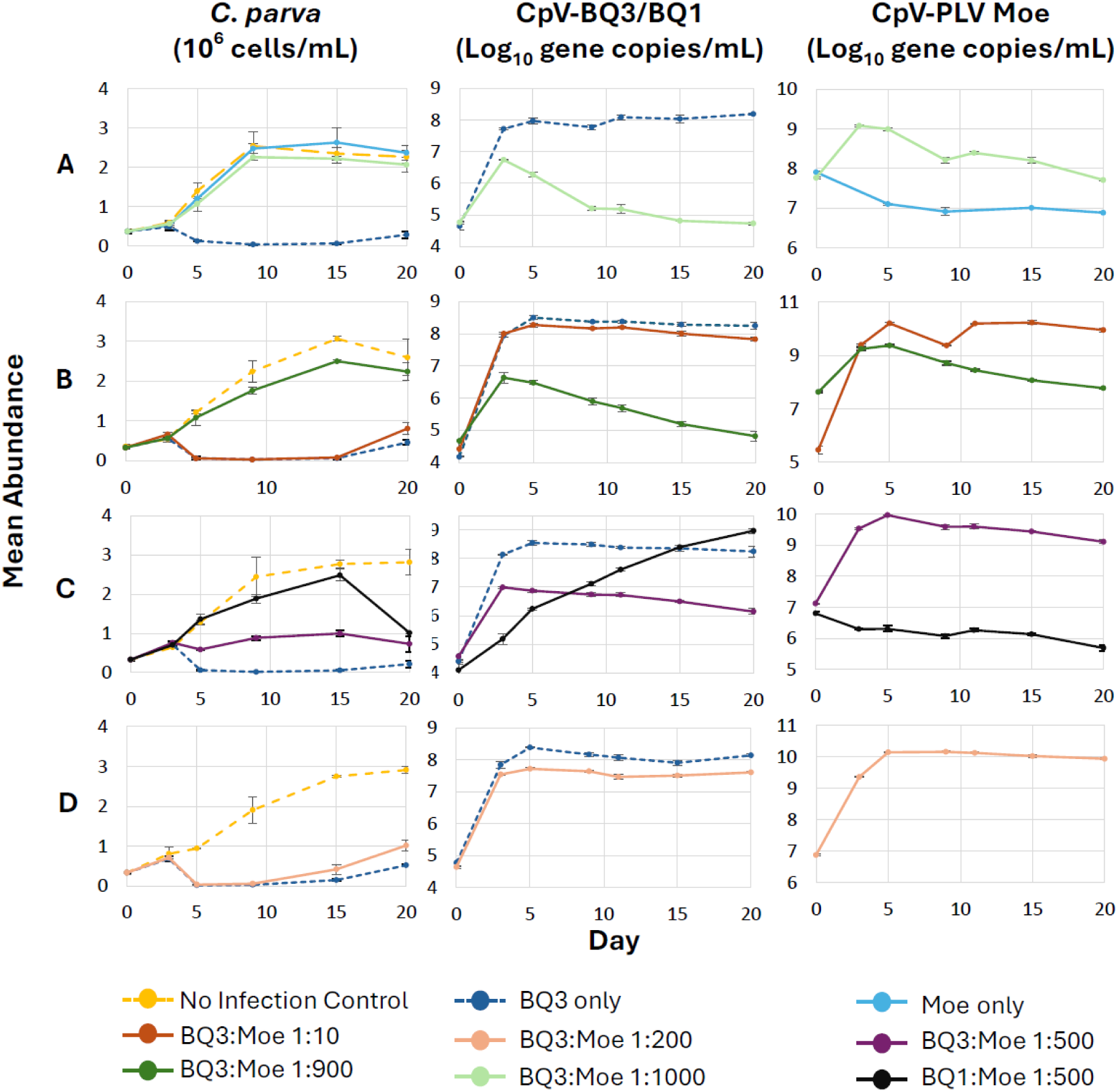
Cell growth and virus replication for all experimental trials. The data presented in columns are labelled at the top of the column; e.g., the first column of panels all present *C. parva* cell abundances. Each row of panels corresponds to a single experimental trial as described in the methods. Error bars represent the standard deviation of replicated experimental incubations.

**Supplementary Figure 4.**
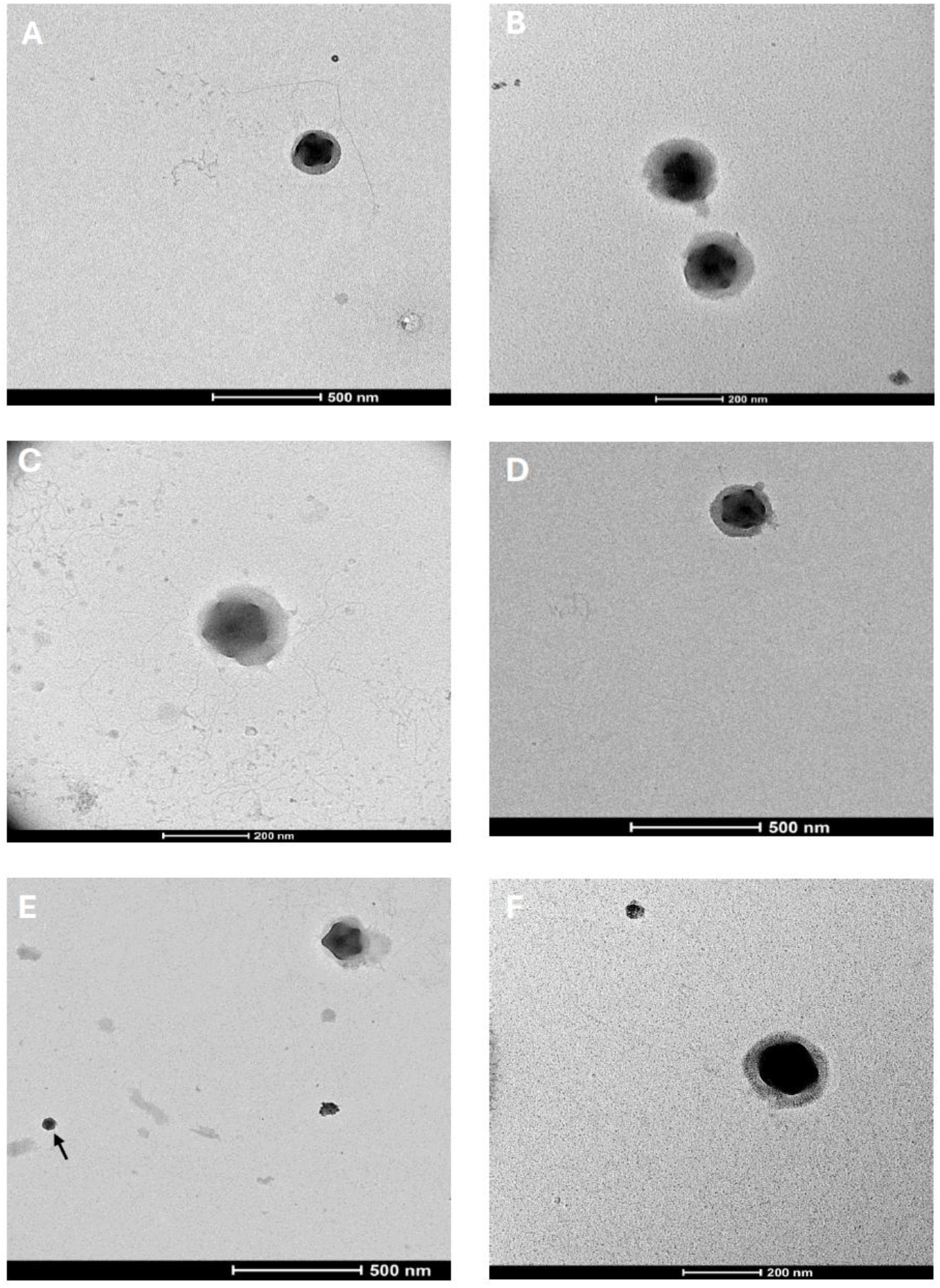
Transmission electron microscopy of viral lysates. (A - D) Micrographs obtained from samples of the 0.45 µm filtration lineage with highly abundant CpV-BQ3 as detected with qPCR. (E & F) Micrographs obtained from samples from the 0.20 µm filtration lineage within which both CpV-BQ3 and CpV-PLV Moe were detected with qPCR. The arrow in panel E points out a particle the same size as Moe.

